# Development of differential sublaminar feedforward inhibitory circuits in CA1 hippocampus requires *Satb2*

**DOI:** 10.1101/2024.01.23.576902

**Authors:** Meretta A. Hanson, Noor Bibi, Alireza Safa, Devipriyanka Nagarajan, Alec H. Marshall, Aidan C. Johantges, Jason C. Wester

## Abstract

Pyramidal cells (PCs) in CA1 hippocampus can be classified by their radial position as deep or superficial and organize into subtype-specific circuits necessary for differential information processing. Specifically, superficial PCs receive fewer inhibitory synapses from parvalbumin (PV)-expressing interneurons than deep PCs, resulting in weaker feedforward inhibition of input from CA3 Schaffer collaterals. Using mice, we investigated mechanisms underlying PC differentiation and the development of this inhibitory circuit motif. We found that expression of the transcriptional regulator SATB2 is biased towards superficial PCs during early postnatal development and necessary to suppress PV+ interneuron synapse formation. In the absence of SATB2, the number of PV+ interneuron synaptic puncta surrounding superficial PCs increases during development to match deep PCs. This results in equivalent inhibitory current strength observed in paired whole-cell recordings, and equivalent feedforward inhibition of Schaffer collateral input. Thus, SATB2 is necessary for superficial PC differentiation and biased feedforward inhibition in CA1.

## INTRODUCTION

Pyramidal cells (PCs) in CA1 provide the output of hippocampal computations to other brain regions. These cells are heterogeneous and include subtypes with unique molecular expression profiles, intrinsic membrane properties, and long-range afferent targets^1–6^. These different subtypes likely engage in overlapping circuits necessary for the diverse tasks performed by the hippocampus^4^. However, CA1 PC subtypes and their functions remain poorly defined, in part because the mechanisms that regulate their differentiation are unknown.

Although diverse, CA1 PCs can be broadly split into two classes based on the radial position of their soma within the stratum pyramidale as deep (closer to the stratum oriens) and superficial (closer to the stratum radiatum)^1^. In mice, deep and superficial PCs exhibit distinct activity patterns during spatial navigation tasks and network oscillations necessary for learning and memory^7–15^. Recent work shows that biases in connectivity and physiology of synapses between inhibitory interneurons and deep versus superficial PCs likely play an important role in their differential activity^16–18^. For example, PV+ basket cells form more perisomatic synapses onto deep than superficial PCs^16,17^, resulting in larger unitary inhibitory synaptic currents^16^. This causes greater feedforward inhibition of CA3 Schaffer collateral input to deep PCs in CA1^19^.Furthermore, superficial PCs target PV+ interneurons for synaptic connections at a higher rate than deep PCs^16^. These synaptic biases likely contribute to differential membrane potential dynamics of deep and superficial PCs during gamma oscillations and sharp-wave ripples^13,17^. However, the mechanisms that guide the formation of these biased circuits during development are unknown.

Clues regarding cell-type-specific circuit development in hippocampal CA1 may be found from prior work on neocortical PC differentiation. In neocortex, a set of crucial transcriptional regulators control PC differentiation during early development^20,21^. One of these is SATB2, a transcription factor and chromatin remodeler that is necessary for differentiation and proper circuit integration of intratelencephalic (IT)-type neocortical PCs^22–26^. In neocortex, *Satb2* expression is necessary to produce IT-type PCs via its repression of *Ctip2*, while *Fezf2* represses *Satb2* and promotes *Ctip2* expression to produce subcerebral-projecting pyramidal tract (PT)-type PCs^22–25,27–33^. Conditional knockout of *Satb2* causes neocortical IT-type PCs to resemble PT-type and disrupts the lamination and synaptic connectivity of interneurons^26,29,33^. In the hippocampus, SATB2 expression is restricted to CA1^34^ and *Satb2* transcription is enriched in superficial PCs^2^. This suggests SATB2 plays a conserved role in CA1 to regulate PC differentiation.

However, the well-characterized laminar expression and interactions of key transcription factors in neocortex do not translate directly to CA1 hippocampus. For example, CTIP2 is preferentially expressed in PCs in deep layers of neocortex^28^ but it is equally expressed in both deep and superficial PCs in CA1 hippocampus^35,36^. Additionally, while *Fezf2* expression is necessary for CTIP2 to be expressed in neocortex, their expression is uncoupled in hippocampus^27^. Thus, it is unknown if transcriptional regulators that determine PC differentiation in neocortex, such as SATB2, play a conserved role in CA1 PC differentiation.

Here, we show that early postnatal expression of SATB2 in CA1 is restricted to superficial PCs and contributes to their differentiation from deep PCs. Importantly, we found that *Satb2* expression in superficial PCs is necessary to suppress PV+ interneuron synapse formation during early postnatal development. This promotes differential feedforward inhibition of CA3 Schaffer collateral input to deep and superficial PCs observed in mature mice. Thus, our data provide insight into the mechanisms underlying PC subtype diversity and the development of biased inhibitory circuits in CA1.

## RESULTS

### SATB2 is preferentially expressed in superficial pyramidal cells in developing CA1 hippocampus

In CA1, there is transcriptomic evidence that *Satb2* is preferentially expressed in superficial PCs of adult mice^2^. However, it is unknown if differential expression of *Satb2* between superficial and deep PCs during early development plays a role in their differentiation. Prior work suggests that SATB2 expression begins later in the hippocampus than neocortex, starting just before birth^34,37^. Thus, we first investigated SATB2 expression in dorsal CA1 at postnatal days (P)0-2 using immunohistochemistry. We used DAPI to identify anatomical landmarks and NeuN to identify developing neurons (**Figure 1Ai**). At P2, we found that PCs that had migrated into the deep half of the hippocampal plate (near the intermediate zone) were mature enough to express NeuN (**Figure 1Aii, carrot**). Because the stratum pyramidale develops “inside-out”, these PCs were likely born during early waves of neurogenesis^38–42^. Strikingly, these early-born deep PCs lacked SATB2 expression (**Figures 1Aiii, carrot**). In contrast, presumed later-born PCs were found throughout the intermediate zone and in the superficial half of the hippocampal plate^39–41,43^. In the intermediate zone, a minority of these immature cells expressed NeuN (**Figure 1Aii, arrow**), but most expressed SATB2 (**Figure 1Aiii, arrow**). Importantly, immature PCs that had migrated to the superficial region of the hippocampal plate also expressed SATB2 (**Figure 1Aiii, arrow outline**). The differential expression pattern of SATB2 and NeuN between deep and superficial cells was consistent from P0 to P2 (**Figures 1A, 1B, and S1**). These data suggest that SATB2 expression in developing CA1 is temporally regulated and preferentially expressed in later-born PCs destined to settle in the superficial half of the stratum pyramidale. SATB2 expression levels increased throughout development, reaching mature levels during the second postnatal week (**Figure 1C**). By P11, immunostaining did not reveal differential expression of SATB2 in deep and superficial PCs (**Figure 1C**).

**Figure 1.**
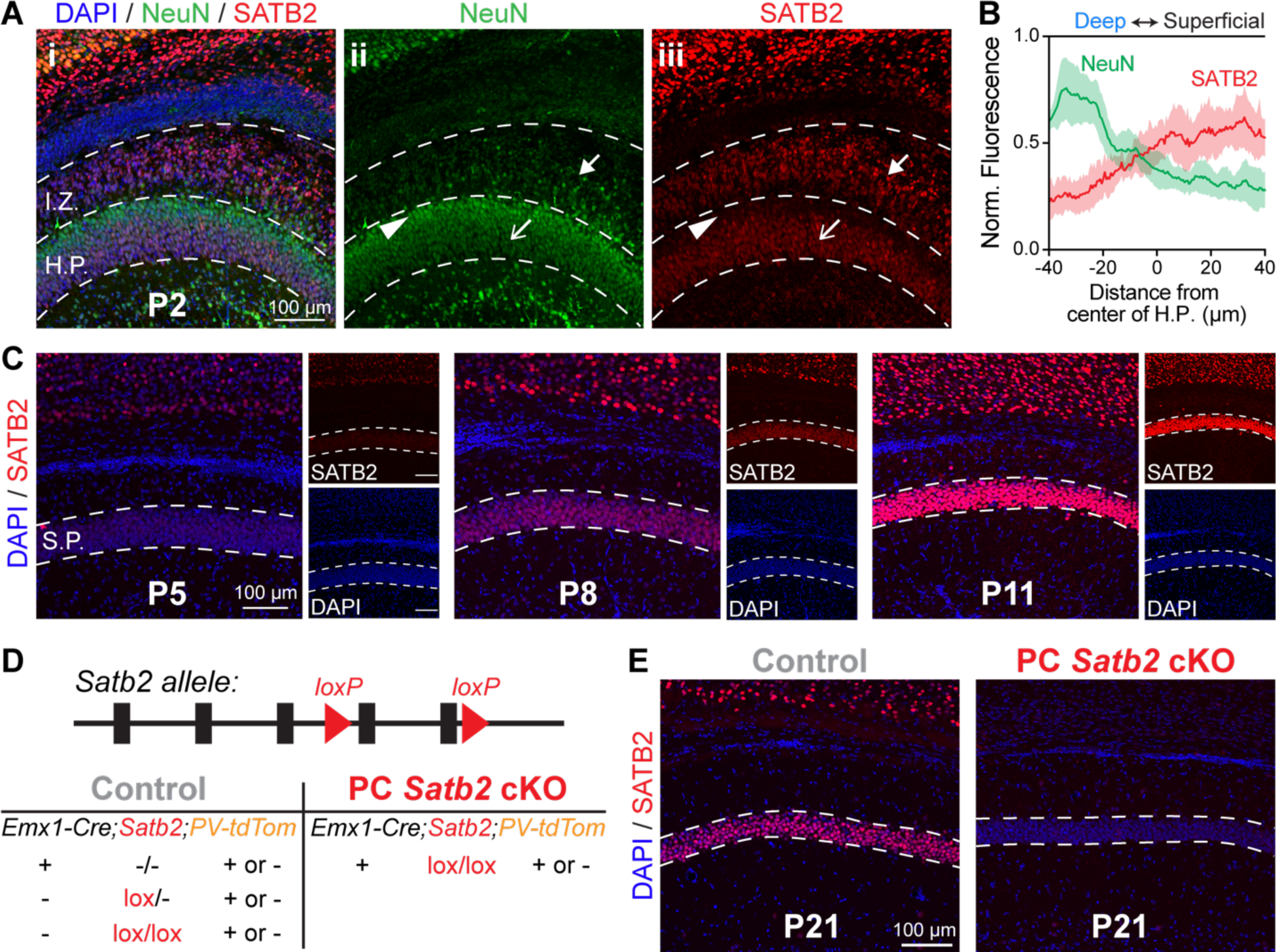
SATB2 is preferentially expressed in superficial PCs and later-born progenitors during early postnatal development. (A) Immunohistochemistry for SATB2 and NeuN at postnatal day 2. HP = hippocampal plate, IZ = intermediate zone. Arrow: SATB2 in migrating cells. Carrot: lack of SATB2 in deep PCs. Arrow outline: SATB2 in superficial PCs. (B) Normalized fluorescence in the HP averaged from three mice at P0, P1, and P2. (C) SATB2 expression in CA1 stratum pyramidale (SP) increases after the first postnatal week. Confocal laser power and gain are the same at each age. (D) Mouse genotypes for control and PC *Satb2* cKO. (E) Verification of PC *Satb2* cKO.

Thus, we hypothesized that SATB2 is necessary for the differentiation of superficial PCs, which in turn may impact their integration into cell-type-specific inhibitory circuits. To test this, we conditionally knocked out (cKO) *Satb2* from PCs during embryonic development by crossing Emx1^IRES-Cre^ mice^44^ to mice with loxP sites flanking exons 4 and 5 of the *Satb2* allele^29,33^ (**Figures 1D and 1E**). We further crossed a subset of these mice to the PV-tdTomato reporter line to target parvalbumin (PV)-expressing inhibitory interneurons^45^ (**Figure 1D**). In brain slices from juvenile mice, we performed whole-cell recordings of deep and superficial PCs and PV+ interneurons in the center of dorsal CA1 hippocampus to investigate cellular and synaptic physiology. We also quantified synapses from PV+ interneurons onto deep and superficial PCs throughout development to provide a structural correlate for differences in inhibition.

### Loss of *Satb2* alters the intrinsic membrane properties of CA1 pyramidal cells

In the neocortex, loss of *Satb2* alters the intrinsic membrane properties of superficial layer (intratelencephalic) projection neurons, causing them to resemble deep layer (pyramidal tract) projection neurons^26,29,46–48^. Thus, we first asked if loss of *Satb2* similarly affects the membrane properties of hippocampal CA1 PCs. Prior work found that the intrinsic membrane properties that differentiate deep and superficial PCs in the center of dorsal CA1 are input resistance and capacitance^5^. Our control data are consistent with those findings, as superficial PCs had a lower input resistance and higher capacitance than deep PCs (**Figures 2A and 2B**). Loss of *Satb2* increased the input resistance and decreased the capacitance of superficial PCs, causing them to resemble deep PCs in control mice, though it was not sufficient to fully reprogram the membrane properties of superficial PCs (**Figures 2A and 2B**). Interestingly, loss of *Satb2* caused voltage sag to increase in both deep and superficial PCs (**Figure 2C**). This suggests *Satb2* regulates H-current channel expression in both PC subpopulations, likely after P8 when *Satb2* is also expressed in deep PCs (**Figure 1C**). Other intrinsic membrane properties were not significantly different in any condition (**Figures 2D-F**). We conclude that *Satb2* expression is necessary but not sufficient for proper development of PC intrinsic membrane properties.

**Figure 2.**
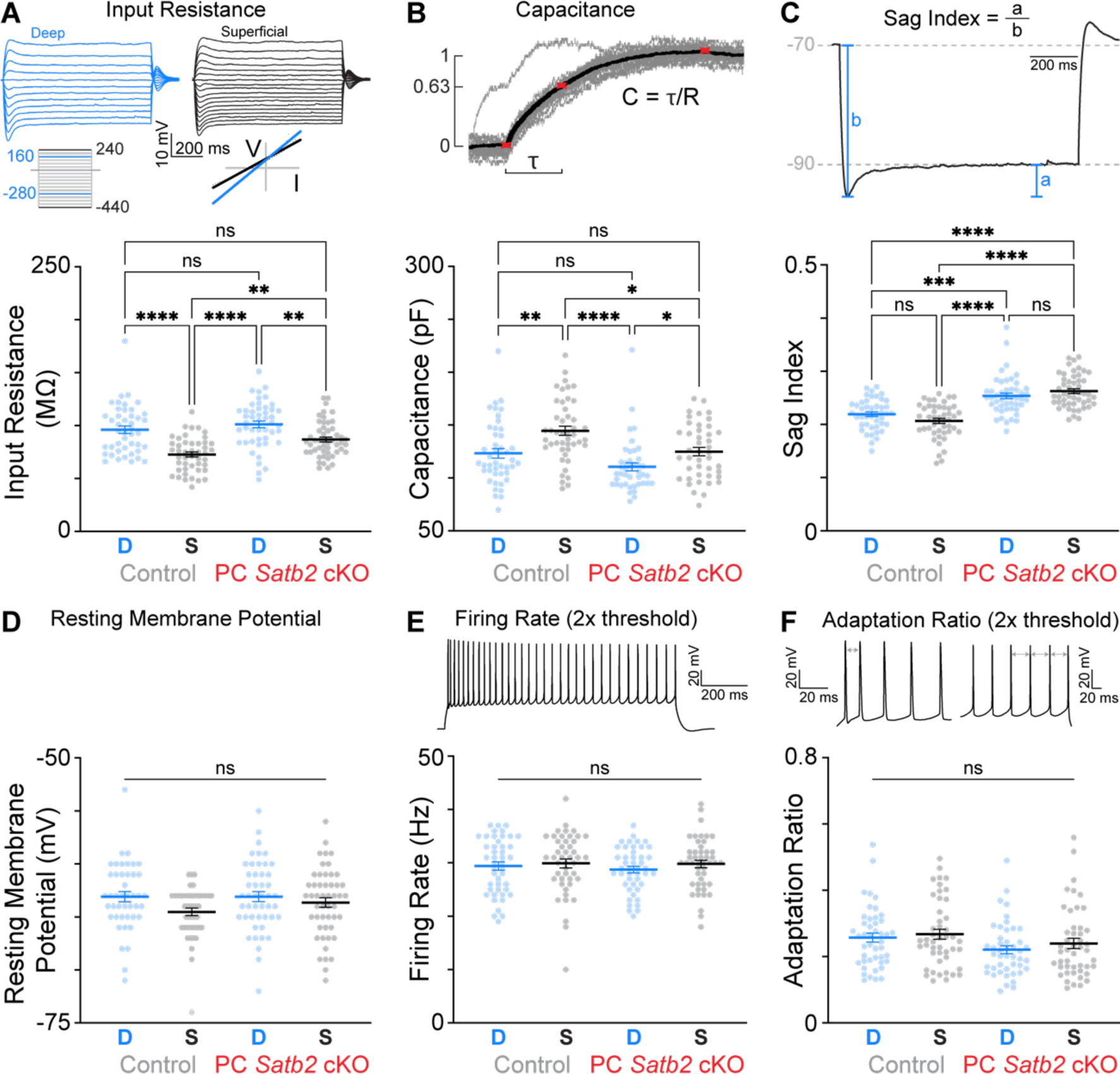
PC *Satb2* cKO alters some intrinsic membrane properties of superficial PCs. (A) Input resistance. Control deep, n = 44 cells; control superficial, n = 46 cells; PC *Satb2* cKO deep, n = 47 cells; PC *Satb2* cKO superficial, n = 50 cells. ** p < 0.01, **** p < 0.0001, Kruskal-Wallis test (H(3) = 44.31, p < 0.0001) with Dunn’s multiple comparisons test. (B) Capacitance. Control deep, n = 44 cells; control superficial, n = 46 cells; PC *Satb2* cKO deep, n = 42 cells; PC *Satb2* cKO superficial, n = 43 cells. * p < 0.05, ** p < 0.01, **** p < 0.0001, Kruskal-Wallis test (H(3) = 32.40, p < 0.0001) with Dunn’s multiple comparisons test. (C) Voltage sag index. Control deep, n = 44 cells; control superficial, n = 46 cells; PC *Satb2* cKO deep, n = 47 cells; PC *Satb2* cKO superficial, n = 50 cells. *** p < 0.001, **** p < 0.0001, Kruskal-Wallis test (H(3) = 69.00, p < 0.0001) with Dunn’s multiple comparisons test. (D) Resting membrane potential. Control deep, n = 44 cells; control superficial, n = 46 cells; PC *Satb2* cKO deep, n = 47 cells; PC *Satb2* cKO superficial, n = 50 cells. Kruskal-Wallis test (H(3) = 6.738, p = 0.0807) with Dunn’s multiple comparisons test. (E) Firing rate. Control deep, n = 45 cells; control superficial, n = 46 cells; PC *Satb2* cKO deep, n = 45 cells; PC *Satb2* cKO superficial, n = 47 cells. Kruskal-Wallis test (H(3) = 2.457, p = 0.4832) with Dunn’s multiple comparisons test. (F) Adaptation ratio. Control deep, n = 45 cells; control superficial, n = 46 cells; PC *Satb2* cKO deep, n = 45 cells; PC *Satb2* cKO superficial, n = 47 cells. Kruskal-Wallis test (H(3) = 6.557, p = 0.0874) with Dunn’s multiple comparisons test. For (A) – (F), data are from 12 control and 14 mutant mice.

### *Satb2* expression is necessary for differential feedforward inhibition of Schaffer collateral input to deep and superficial pyramidal cells

Next, we investigated if *Satb2* expression is necessary for the development of PC cell-type-specific circuits in CA1. Previously, Masurkar *et al.* ^19^ found that input from CA3 Schaffer collateral projections evoke weaker depolarization of deep than superficial PCs. This is due to enhanced feedforward inhibition of deep PCs, rather than differences in the strength of excitatory Schaffer collateral synapses^19^. We replicated these experiments in control and PC *Satb2* cKO mice to determine if *Satb2* is necessary for proper development of this circuitry. We placed a stimulating electrode in stratum radiatum and performed whole-cell recordings of deep and superficial pyramidal cells while increasing stimulus intensity (**Figure 3A**). We measured responses in current-clamp with the resting membrane potential biased to –70 mV and then switched to voltage-clamp with a holding potential of –70 mV. This allowed us to confirm that differences in evoked membrane potential were due to underlying synaptic currents, rather than differences in intrinsic membrane properties between deep and superficial PCs, such as input resistance (**Figure 2A)**. First, we stimulated afferents in the presence of GABA_A_ and GABA_B_ receptor blockers to assay excitatory synapse strength (**Figure 3A**). Consistent with previous work^19^, we found that evoked excitatory postsynaptic potentials (EPSPs) were of comparable amplitude in deep and superficial PCs (**Figure 3Bi, top**). Furthermore, we confirmed this was due to comparable amplitude of evoked excitatory postsynaptic currents (EPSCs) (**Figure 3Bi, bottom**). PC *Satb2* cKO did not alter the strength of these afferent inputs onto deep or superficial PCs (**Figure 3Bii**).

**Figure 3.**
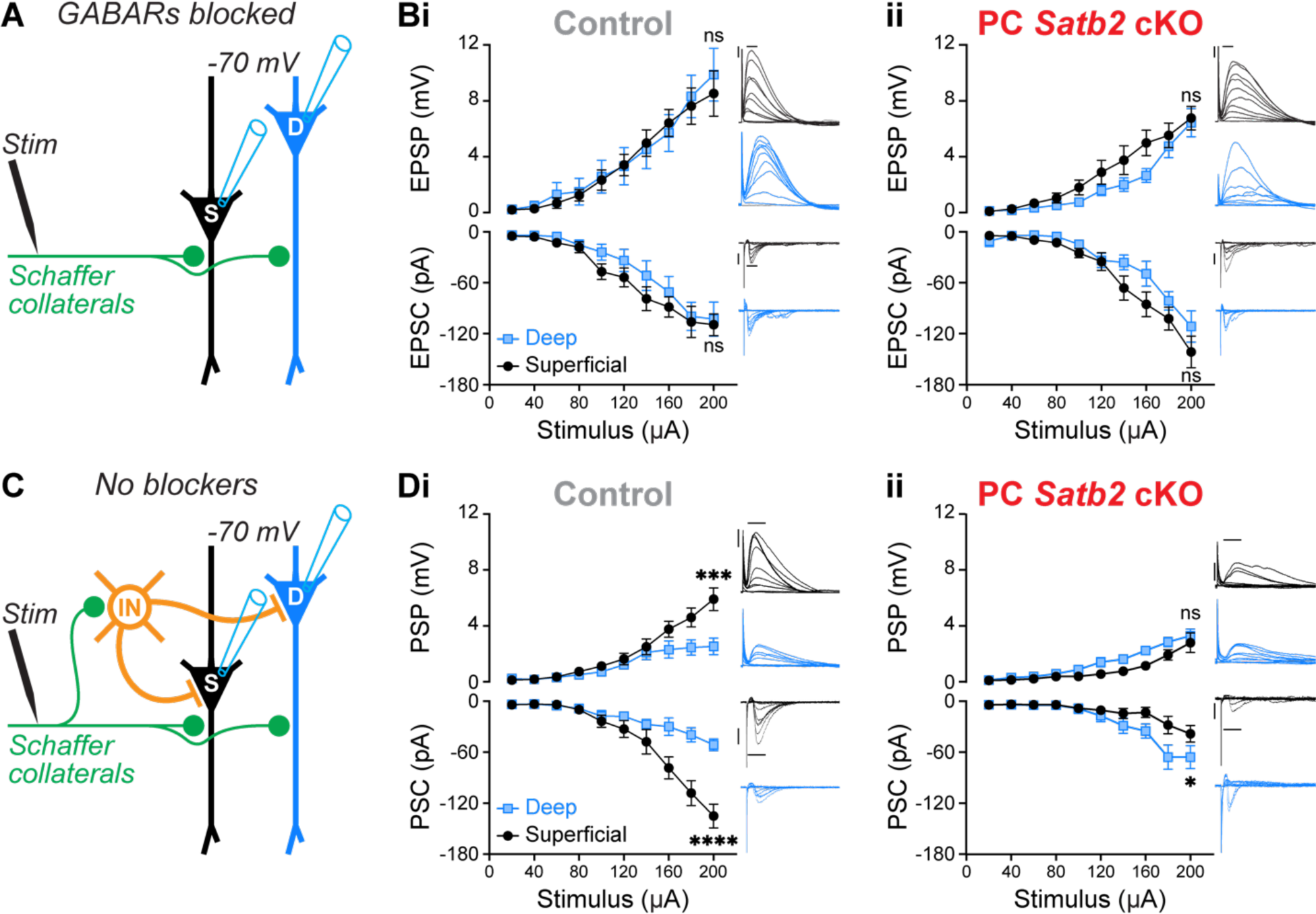
PC *Satb2* cKO increases Schaffer collateral-driven feedforward inhibition of superficial PCs to match deep PCs. (A) Experimental configuration with inhibition blocked. (B) Input-output curves of PC synaptic responses to Schaffer collateral stimulation with inhibition blocked. PCs recorded in current-clamp (top) and voltage-clamp (bottom). (i) Control. Two-way ANOVA of EPSP (F[9, 137] = 0.1533, p = 0.99) and EPSC (F[9, 140] = 0.3397, p = 0.96). n = 8 deep and 8 superficial PCs from 8 mice. (ii) PC *Satb2* cKO. Two-way ANOVA of EPSP (F[9, 150] = 0.8709, p = 0.55) and EPSC (F[9, 150] = 1.088, p = 0.37). n = 9 deep and 8 superficial PCs from 4 mice. Insets are example responses from single cells. Scale bars are 4 mV, 100 pA, 40 ms. (C) Experimental configuration with inhibition intact. (D) Same as (B) with inhibition intact. (i) Control. Two-way ANOVA of PSP (F[9, 150] = 3.631, *** p = 0.0004) and PSC (F[9, 150] = 7.157, **** p < 0.0001). n = 8 deep and 9 superficial PCs from 8 mice. (ii) PC *Satb2* cKO. Two-way ANOVA of PSP (F[9, 160] = 0.9862, p = 0.45) and PSC (F[9, 160] = 2.035, * p = 0.04). n = 9 deep and 9 superficial PCs from 6 mice. Insets are example responses from single cells. Scale bars same as (B).

Next, we repeated these experiments without GABA receptor blockers to determine if *Satb2* regulates feedforward inhibition of these afferents (**Figure 3C**). In control mice, we found that the amplitudes of evoked postsynaptic potentials (PSPs) were smaller in deep relative to superficial PCs as stimulus intensity increased (**Figure 3Di, top**), consistent with Masurkar *et al.* ^19^. This was due to smaller evoked synaptic inward postsynaptic currents (PSCs) in deep PCs (**Figure 3Di, bottom**), which is consistent with greater recruitment of shunting feedforward inhibition. Strikingly, in PC *Satb2* cKO mice, feedforward inhibition of superficial PCs increased to match that observed in deep PCs (**Figure 3Dii**). Furthermore, the amplitudes of PSPs and PSCs were not different between deep PCs in control and PC *Satb2* cKO mice. Thus, these data show that expression of *Satb2* in superficial, but not deep PCs, is crucial for the development of differential feedforward inhibition of Schaffer collateral input.

Finally, we investigated if these findings extended to other afferent pathways into CA1. We repeated the above experiments but placed the stimulating electrode in the stratum lacunosum moleculare to target input from the entorhinal cortex and thalamus (**Figure S2**). We found no differences in the amplitudes of evoked responses between deep and superficial PCs in either control or PC *Satb2* cKO mice, both with and without GABA receptor blockers (**Figure S2**). We conclude that the role of *Satb2* in the development of feedforward inhibitory circuits is selective for interneurons involved in Schaffer collateral projections to superficial PCs.

### *Satb2* expression decreases the strength of unitary synaptic connections from PV+ interneurons to superficial pyramidal cells by regulating synapse number

Previously, Lee *et al.* ^16^ used paired whole-cell recordings to show that synaptic connections from PV+ basket cells to PCs in CA1 evoke larger inhibitory currents in deep than superficial cells. This is likely the primary mechanism underlying the differential strength of feedforward inhibition in CA1^19^ (**Figure 3Di**). Thus, we used the PV-tdTomato reporter line^45^ (**Figure 1D**) to investigate the connectivity and physiology of synapses between PCs and PV+ interneurons using paired whole-cell recordings in control and PC *Satb2* cKO conditions. In control mice, we confirmed previous findings^16^ that the connectivity rate of inhibitory connections from PV+ interneurons to PCs is comparable between deep and superficial PCs, but the average amplitude of inhibitory postsynaptic currents (IPSCs) is larger in deep PCs (**Figure 4A**). Furthermore, we confirmed that this differential IPSC amplitude is determined by PC subtype identity. We patched a single PV+ interneuron and then sequentially patched four neighboring PCs, alternating between deep and superficial (**Figure S3A**). This PV+ interneuron provided a synaptic connection to all four recorded PCs. Strikingly, IPSCs were larger in both deep PCs relative to either superficial PC, demonstrating that the identity of the postsynaptic PC regulates the strength of this synapse (**Figure S3B**).

Next, we investigated if *Satb2* expression is necessary for the bias in unitary synaptic connection strength between PV+ interneurons and PCs. Before performing paired whole-cell recordings, we first determined if PC *Satb2* cKO causes non-cell-autonomous changes in PV+ interneuron development using immunostaining for PV (**Figure S4A**). Interestingly, we found that the radial positioning of PV+ interneurons was disrupted in juvenile PC *Satb2* cKO mice: more PV+ interneurons were found in stratum oriens at the expense of stratum pyramidale (**Figure S4B**). This was due to a migration deficit, not loss of cells, because the total number of PV+ interneurons in CA1 was unaffected (**Figure S4C**). Thus, like in neocortex, loss of *Satb2* in PCs alters the radial position of interneurons in CA1^26,31,49,50^. However, PV+ interneurons recorded in the stratum pyramidale exhibited normal morphology (**Figure S4D**) and intrinsic membrane properties (**Figures S4E-J**); although we noted a decrease in average input resistance (**Figure S4F**). Thus, to investigate synaptic physiology in PC *Satb2* cKO mice, we restricted our whole-cell recordings of PV+ interneurons to those that successfully migrated to the stratum pyramidale.

In PC *Satb2* cKO mice, we found that synaptic connectivity rates between PCs and PV+ interneurons were comparable to control (**Figure 4B, top**). However, the average amplitude of IPSCs recorded in superficial PCs increased to match that observed in deep PCs (**Figure 4B, bottom**). For each synaptic connection, we delivered ten trials of presynaptic action potential trains. This allowed us to quantify synaptic potency, which only considers trials in which a presynaptic action potential evoked an IPSC, and synaptic failure rate, which is the fraction of trials that failed to evoke an IPSC^51^. These measurements provide insight into possible pre– and postsynaptic mechanisms underlying differences in IPSC amplitude. In control mice, the average IPSC potency was significantly greater in deep than superficial PCs; strikingly, in PC *Satb2* cKO mice, IPSC potency was identical in both subtypes (**Figure 4C**). To determine if changes at the synapses onto both deep and superficial PCs contributed to this finding, we replotted the data to directly compare each subtype between conditions (**Figure 4D**). Importantly, the average IPSC potency observed in deep PCs was not different between conditions (**Figure 4D, left**), but in superficial PCs it was significantly greater in *Satb2* cKO mice (**Figure 4D, right**). Thus, *Satb2* expression in superficial PCs, but not deep, is necessary for the development of differential inhibitory synapse strength. Finally, to determine if presynaptic mechanisms contribute to differences in average IPSC amplitude, we compared synaptic failure rates^52^. We found no differences between deep and superficial PCs in either condition (**Figure 4E**).

These data suggest that *Satb2* expression regulates either the number of synapses made by each PV+ interneuron or postsynaptic receptor density. Previous work found that a greater number of PV+ synaptic puncta surround deep than superficial PCs^16,17^. Thus, we next performed immunohistochemistry for synaptogamin-2 (SYT2) which labels presynaptic puncta from PV+ interneurons^53^. At postnatal day 21, SYT2 expression in CA1 was concentrated in the stratum pyramidale and was not altered by PC *Satb2* cKO (**Figures 4F and 4G**). Thus, localization of PV+ interneuron synapses to stratum pyramidale was not grossly disrupted by PC *Satb2* cKO. We next quantified the density of SYT2+ puncta surrounding the somas of individual deep and superficial PCs. In control mice, PV+ interneurons formed significantly more synapses onto deep than superficial PCs, consistent with previous findings^16^ (**Figures 4H and 4I**). Strikingly, in PC *Satb2* cKO mice, the number of SYT2+ puncta surrounding superficial PCs increased relative to control to match the number of puncta surrounding deep PCs in both conditions (**Figures 4H and 4I**). Together, these data show that *Satb2* expression in superficial PCs negatively regulates the strength of inhibitory input from PV+ interneurons by reducing the number of inhibitory synapses. This is necessary to establish differential feedforward inhibition of Schaffer collateral input to deep and superficial PCs in CA1 (**Figure 3**).

### *Satb2* expression in superficial pyramidal cells suppresses the formation of PV+ interneuron synapses during early postnatal development

Next, we investigated the developmental mechanism by which *Satb2* expression regulates the number of inhibitory synapses surrounding superficial PCs in CA1. We hypothesized that *Satb2* expression could either suppress synapse formation or promote elimination of excess synapses prior to postnatal day 21. In both CA1 and neocortex, perisomatic inhibitory synapse formation increases rapidly around postnatal day 10^54–56^. Thus, we next quantified SYT2+ puncta surrounding the somas of deep and superficial PCs at postnatal days 10 and 14 in control and PC *Satb2* cKO mice. Although PV expression is low before the second postnatal week^57,58^, SYT2 is still a reliable marker for future PV+ presynaptic puncta (**Figure 5A**). At postnatal day 10, during early PV+ interneuron synaptogenesis, we found a clear bias in the number of puncta surrounding deep PCs relative to superficial PCs in control mice (**Figure 5B**). Thus, differences in synapse formation, not synapse elimination, underly the development of differential inhibitory synaptic strength. Interestingly, this bias was also apparent in PC *Satb2* cKO mice; however, relative to controls, the number of puncta around deep PCs was reduced and the number of puncta around superficial PCs was increased (**Figure 5B**). By postnatal day 14, the ratios of SYT2+ puncta around deep and superficial PCs in both conditions resembled those observed at postnatal day 21 (**Figures 4I, 5C, and 5D**). In particular, the number of SYT2+ synapses around deep and superficial PCs increased in PC *Satb2* cKO mice to match that observed for deep PCs in controls (**Figures 5C and 5D**).

**Figure 4.**
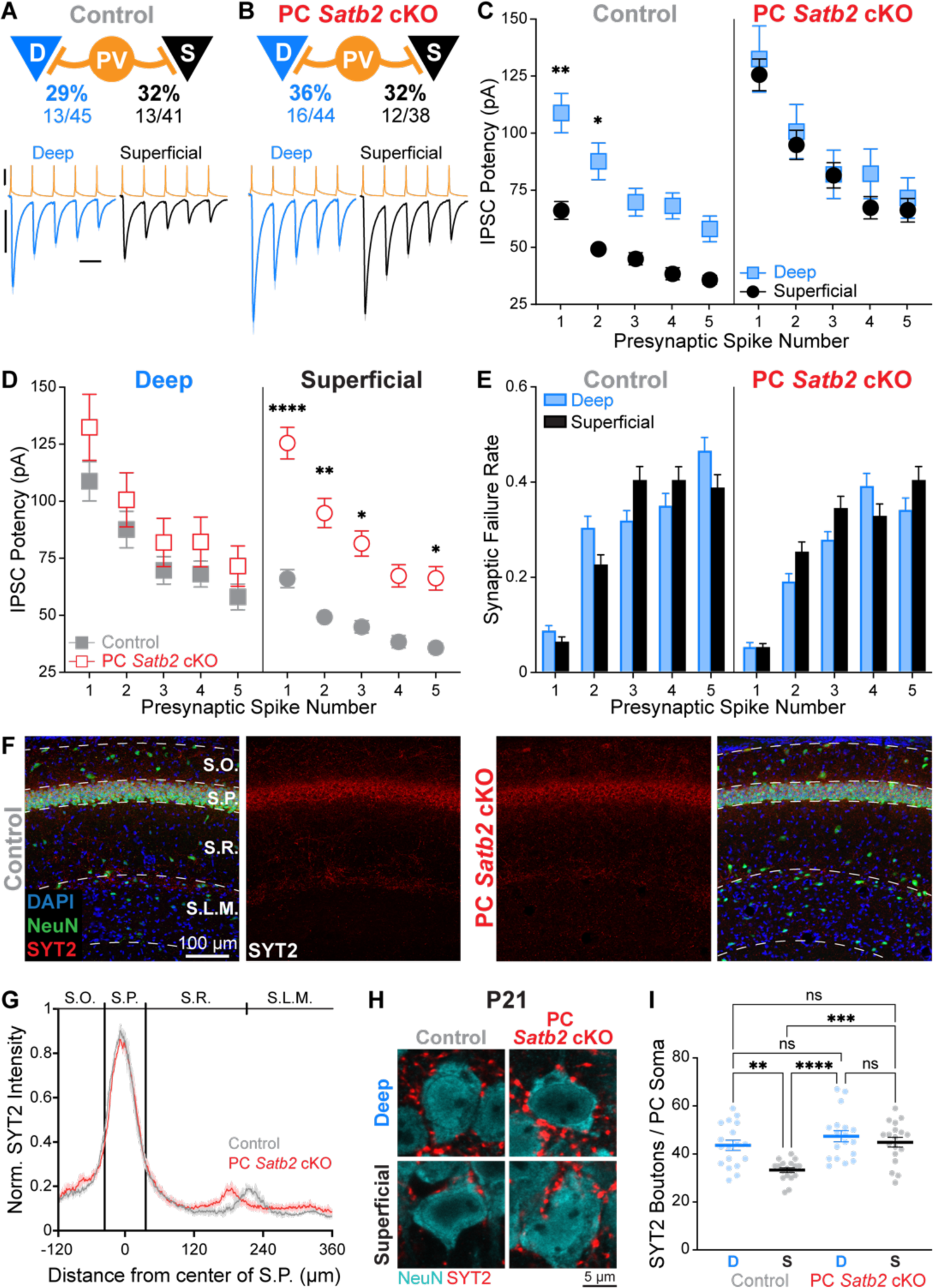
PC *Satb2* cKO increases PV+ interneuron inhibition of superficial PCs to match deep PCs. (A) Connectivity rates and physiology of synapses from PV+ interneurons to deep and superficial PCs in control mice. Shown are the average amplitudes (± SEM) of all postsynaptic responses from connected pairs. n = 13 deep and 13 superficial pairs, 10 trials per pair, from 13 mice. Scale bars are 40 mV, 40 pA, 20 ms. (B) Same as (A) for PC *Satb2* cKO mice. n = 16 deep pairs 12 superficial pairs, 10 trials per pair, from 12 mice. Scale same as (A). (C) Average IPSC potency (synaptic failures removed) as a function of PV + interneuron presynaptic spike number. n’s for IPSCs averaged for spikes 1 – 5: control deep = 119, 91, 89, 85, 70; control superficial = 122, 101, 78, 78, 80; PC *Satb2* cKO deep = 152, 130, 116, 98, 106; PC *Satb2* cKO superficial = 114, 90, 79, 81, 72. n’s for mice and synaptic connections for each condition are the same as (A) and (B). * p < 0.05, ** p < 0.01, two-way ANOVA (F[3, 1918] = 26.67, p < 0.0001) with Tukey’s multiple comparisons test. (D) Same as (C) replotted to show that only synapses onto superficial PCs are enhanced in PC *Satb2* cKO mice. * p < 0.05, ** p < 0.01, **** p < 0.0001, two-way ANOVA with Tukey’s multiple comparisons test same as (C). (E) Average synaptic failure rate as a function of PV+ interneuron presynaptic spike number. n’s for each presynaptic spike: control deep = 130; control superficial = 130; PC *Satb2* cKO deep = 160; PC *Satb2* cKO superficial = 120. n’s for mice and synaptic connections for each condition are the same as (A) and (B). Post hoc analysis revealed no significant differences between individual groups, two-way ANOVA (F[3, 2680] = 6.035, p = 0.0004) with Tukey’s multiple comparisons test. (F) The localization of SYT2 expression to the stratum pyramidale is not altered by PC *Satb2* cKO. (G) Quantification of SYT2 localization (as fluorescence intensity) across CA1 strata in control and PC *Satb2* cKO (n = 2 images from 2 mice per condition). (H) SYT2+ presynaptic puncta surrounding deep and superficial PCs in control and PC *Satb2* cKO at P21. Note fewer puncta around superficial PCs in control mice. (I) Quantification of SYT2+ puncta surrounding deep and superficial PCs in control and PC *Satb2* cKO at P21. Note that in PC *Satb2* cKO mice the number of puncta surrounding superficial PCs increased to match that observed in deep PCs (n = 6 deep and 6 superficial cells per mouse, 3 mice per condition). ** p < 0.01, *** p < 0.001, **** p < 0.0001, one-way ANOVA (F[3, 68] = 10.27, p < 0.0001) with Tukey’s multiple comparisons test.

**Figure 5.**
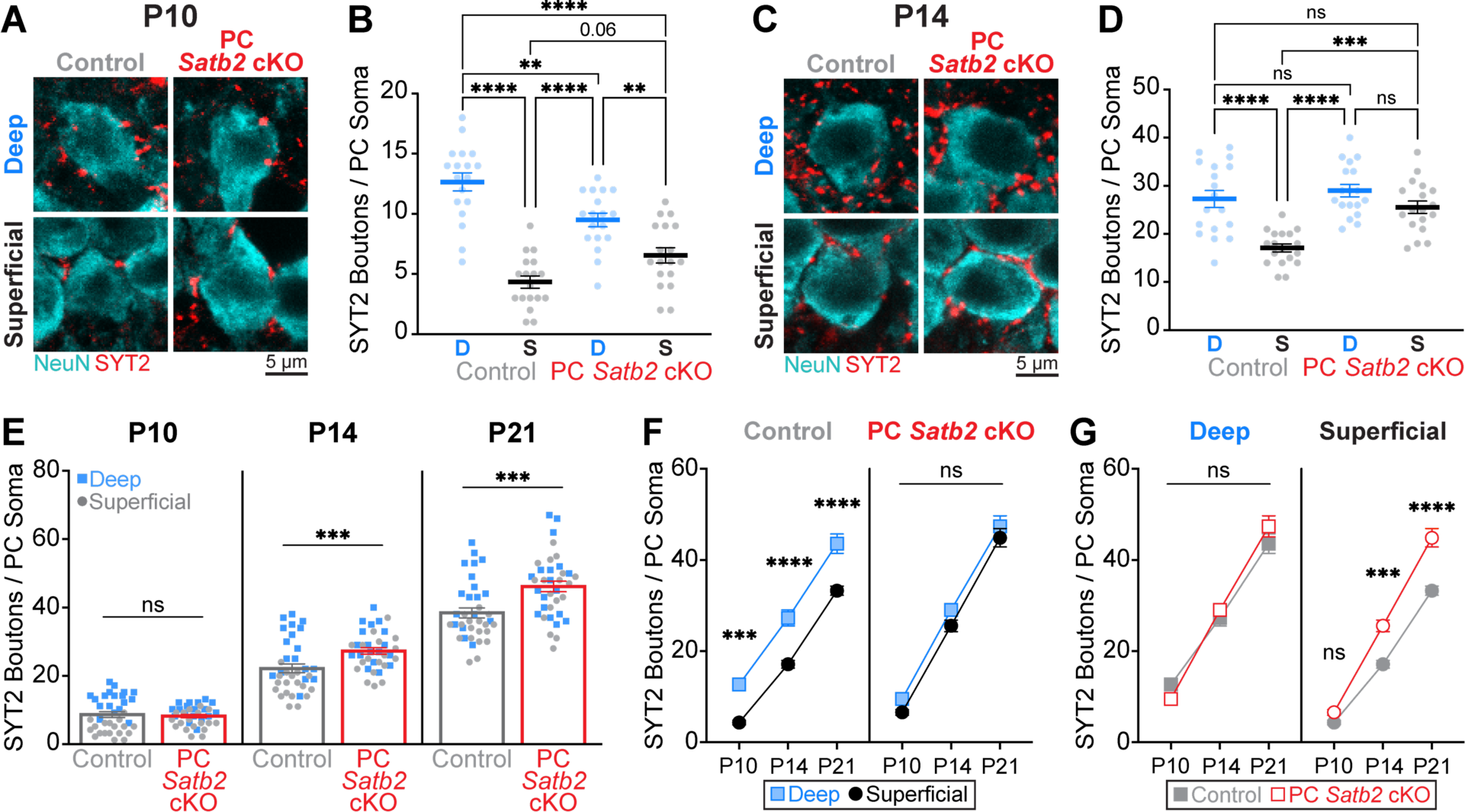
PC *Satb2* cKO increases number of PV+ interneuron synapses onto superficial PCs to match deep PCs. (A) SYT2+ presynaptic puncta surrounding deep and superficial PCs in control and PC *Satb2* cKO at P10. Note fewer puncta around superficial PCs in control mice. (B) Quantification of SYT2+ presynaptic puncta at P10. Note strong inhibitory bias is present during early development in control mice and weakened in PC *Satb2* cKO mice (n = 6 deep and 6 superficial cells per mouse, 3 mice per condition). ** p < 0.01, **** p < 0.0001, one-way ANOVA (F[3, 68] = 33.89, p < 0.0001) with Tukey’s multiple comparisons test. (C) SYT2+ presynaptic puncta surrounding deep and superficial PCs in control and PC *Satb2* cKO at P14. Note the fewest SYT2+ puncta surround control superficial PCs. (D) Quantification of SYT2+ presynaptic puncta at P14. Note that in PC *Satb2* cKO mice the number of puncta surrounding superficial PCs increased to match that observed in both control and mutant deep PCs (n = 6 deep and 6 superficial cells per mouse, 3 mice per condition). *** p < 0.001, **** p < 0.0001, one-way ANOVA (F[3, 68] = 15.85, p < 0.0001) with Tukey’s multiple comparisons test. (E) SYT2+ presynaptic puncta surrounding deep and superficial PCs pooled to compare total number of puncta between control and PC *Satb2* cKO mice at each age. Note that total STY2+ puncta are equivalent at P10 but greater in PC *Satb2* cKO mice by P14. Furthermore, at P10 it is already clear that the distributions of SYT2+ puncta numbers diverge between deep and superficial PCs in control mice but overlap in PC *Satb2* cKO mice. Mann-Whitney U test at P10 (U = 633.5, p = 0.8729), P14 (U = 356, *** p = 0.0008), and P21 (U = 341.5, *** p = 0.0004). (F) Average number of SYT2+ presynaptic puncta surrounding each PC subtype in control and PC *Satb2* cKO mice as a function of age. Note that in deep and superficial PCs, STY2+ puncta densities are equivalent at each age in PC *Satb2* cKO mice. *** p < 0.001, **** p < 0.0001, two-way ANOVA (F[3, 204] = 34.29, p < 0.0001) with Tukey’s multiple comparisons test. (G) Same as (F) replotted to show that only SYT+ puncta surrounding superficial PCs increase in number in PC *Satb2* cKO mice. *** p < 0.001, **** p < 0.0001, two-way ANOVA with Tukey’s multiple comparisons test same as (F).

These data suggest that *Satb2* expression is necessary to suppress synapse formation onto superficial PCs. However, it was unclear why we observed fewer SYT2+ puncta around deep PCs and only a partial increase around superficial PCs at postnatal day 10 in PC *Satb2* cKO mice (**Figure 5B**). We hypothesized that at postnatal day 10, the rate of inhibitory synapse formation is limited. Thus, in the absence of *Satb2* expression, excess synapse formation onto superficial PCs occurred at the expense of synapse formation onto deep PCs. We compared the average total number of SYT2+ puncta (deep and superficial pooled) between control and PC *Satb2* cKO mice at postnatal days 10, 14, and 21 (**Figure 5E**). At postnatal day 10, the average number of SYT2+ synapses were not different between control and PC *Satb2* cKO mice (**Figure 5E, left**). In contrast, by postnatal day 14 there were more total SYT2+ synapses in PC *Satb2* cKO mice (**Figure 5E, middle**). Importantly, this coincided with the increase in SYT2+ puncta around deep and superficial PCs in PC *Satb2* cKO mice to match deep PCs in control (**Figure 5D**). Thus, at postnatal day 10, superficial PCs in PC *Satb2* cKO mice recruit PV+ synapses at the expense of deep PCs. However, by postnatal day 14, the rate of synaptogenesis is sufficient for PV+ interneurons to distribute typical densities to deep PCs and excessive densities to superficial PCs in PC *Satb2* cKO mice.

Finally, we compared the average number of SYT2+ puncta between control and PC *Satb2* cKO mice and between deep and superficial PCs as a function of age (**Figures 5F and 5G**). This confirmed that the number of SYT2+ puncta surrounding deep cells exceeds that surrounding superficial PCs in control mice but not PC *Satb2* cKO mice throughout postnatal development (**Figure 5F**). Furthermore, this is due to excessive synapse formation selectively onto superficial PCs in *Satb2* cKO mice, which is apparent by postnatal day 14 (**Figure 5G**).

In summary, our data show that preferential feedforward inhibition of Schaffer collateral input to deep PCs observed in mature mice is hardwired during early development. To establish this circuit motif, early postnatal expression of *Satb2* is necessary to suppress the formation of PV+ interneuron synapses onto superficial PCs.

### *Satb2* does not regulate differential pyramidal cell excitation of PV+ interneurons

Finally, CA1 PCs provide excitatory synaptic input to PV+ interneurons to form a recurrent excitatory-inhibitory loop^59^. Lee *et al.* ^16^ observed in paired recordings that the connectivity rate from superficial PCs to PV+ interneurons is higher than from deep PCs. Thus, we investigated if *Satb2* also regulates the development of this differential connectivity. Consistent with Lee *et al.* ^16^, we found a bias in the probability of observing synaptic connections from superficial PCs to PV+ interneurons (**Figure 6A, top**). However, this bias was not affected in PC *Satb2* cKO mice and connectivity rates were comparable to control (**Figure 6A, bottom**). Interestingly, we also observed larger average excitatory postsynaptic current (EPSC) amplitude in connections from superficial PCs (**Figure 6B**), which was also unaffected in PC *Satb2* cKO mice (**Figure 6C**). This bias in average EPSC amplitude was primarily due to enhanced EPSC potency of superficial connections (**Figure 6D**). However, we also noted greater presynaptic release probability from superficial PCs (**Figure 6E**). Together, these data extend previous findings to show that deep and superficial PCs engage in distinct, yet connected, excitatory-inhibitory loops in CA1: superficial PCs provide biased excitation to PV+ interneurons, which in turn provide biased inhibition to deep PCs (**Figure 7A**). Expression of *Satb2* in superficial PCs is crucial for the proper development of inhibitory, but not excitatory, synapses within this loop (**Figure 7B**).

**Figure 6.**
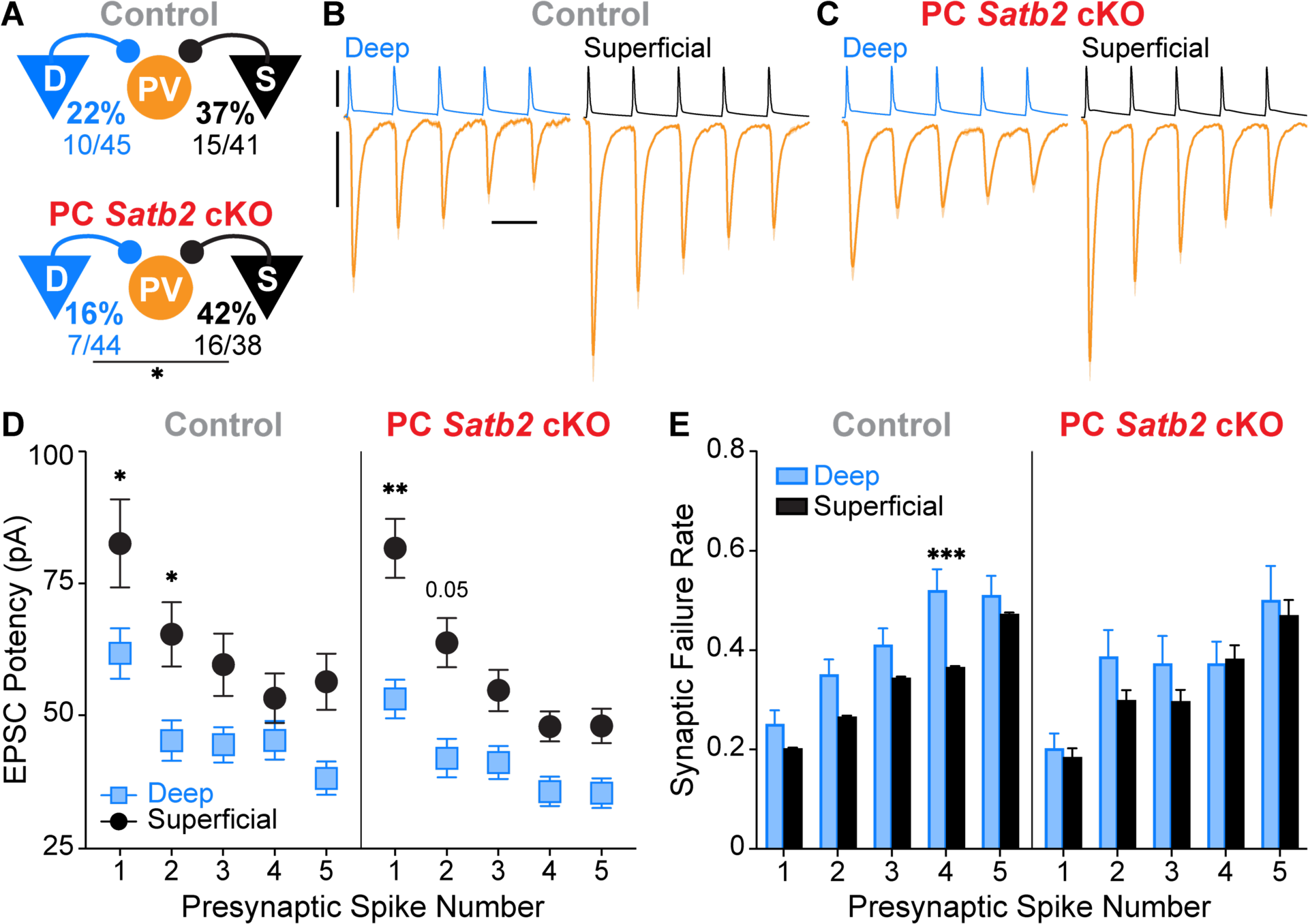
Superficial PCs provide biased synaptic input to PV+ interneurons that is not affected by PC *Satb2* cKO. (A) Synaptic connectivity rates in paired recordings between PCs and PV+ interneurons in control and PC *Satb2* cKO mice. * p < 0.05, Fisher’s exact test. (B) Average (± SEM) amplitudes of all postsynaptic responses from connected pairs in control mice. n = 10 deep and 15 superficial pairs, 10 trials per pair, from 13 mice. Scale bars are 80 mV, 20 pA, 20 ms. (C) Same as (B) for *Satb2* cKO mice. n = 7 deep and 16 superficial pairs, 10 trials per pair, from 12 mice. (D) Average EPSC potency (synaptic failures removed) as a function of PC presynaptic spike number. n’s for EPSCs averaged for spikes 1 – 5: control deep = 75, 65, 59, 48, 49; control superficial = 102, 93, 83, 80, 66; PC *Satb2* cKO deep = 56, 43, 44, 44, 35; PC *Satb2* cKO superficial = 116, 101, 99, 83, 77. n’s for mice and synaptic connections for each condition are the same as (B) and (C). * p < 0.05, two-way ANOVA (F[3, 1489] = 14.91, p < 0.0001) with Tukey’s multiple comparisons test. (E) Average synaptic failure rate as a function of PC presynaptic spike number. n’s for each presynaptic spike: control deep n = 100, control superficial n = 150, mutant deep n = 70, and mutant superficial n = 160. n’s for mice and synaptic connections for each condition are the same as (B) and (C). *** p < 0.001, two-way ANOVA (F[3, 2330] = 9.754, p < 0.0001) with Tukey’s multiple comparisons test.

**Figure 7.**
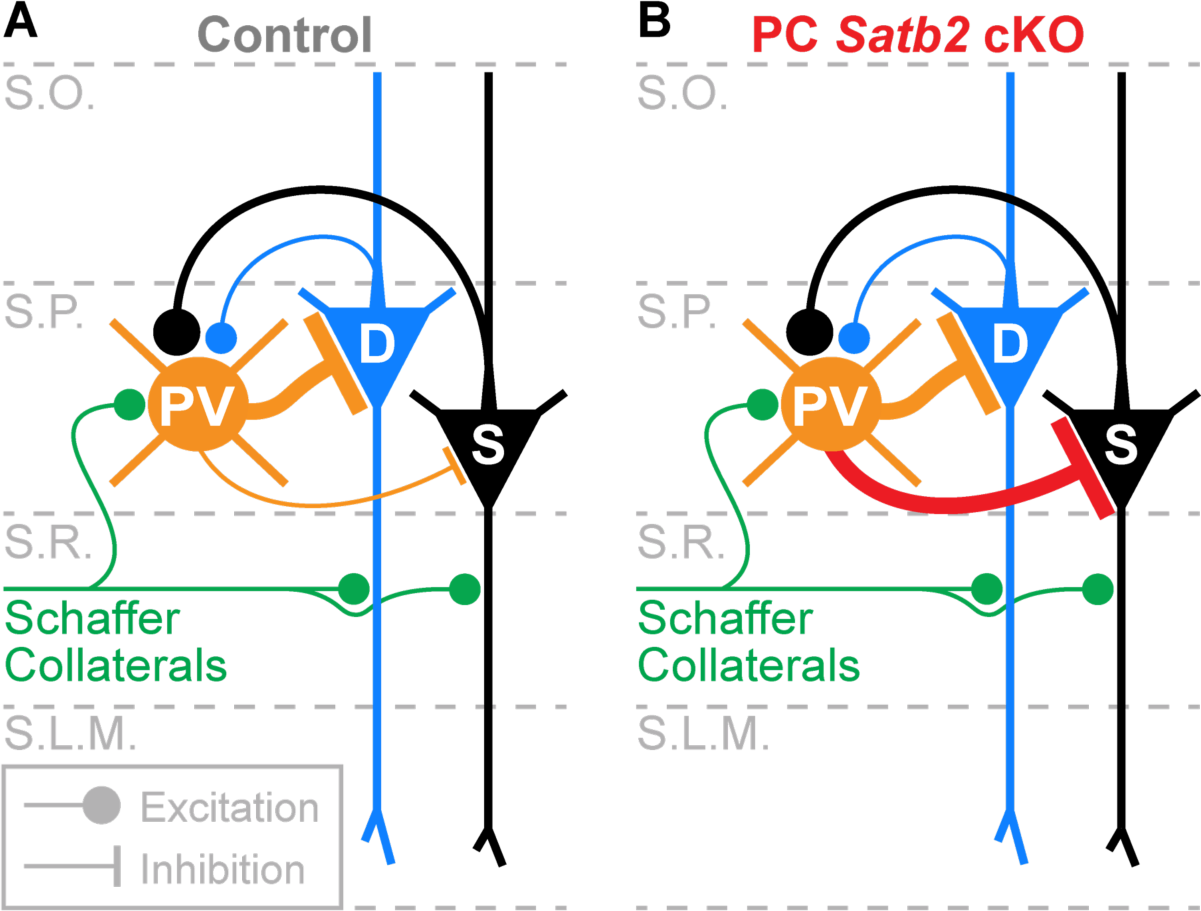
SATB2 is necessary to establish the excitatory-inhibitory loop between deep and superficial PCs in CA1 hippocampus. (A) Superficial PCs provide biased excitation to local PV+ interneurons which, in turn, preferentially inhibit deep PCs. (B) Conditional knockout of *Satb2* from PCs results in equal inhibition of deep and superficial PCs by PV+ interneurons. PC *Satb2* cKO does not impact preferential excitation of PV+ interneurons by superficial PCs.

## DISCUSSION

Pyramidal cells in CA1 hippocampus are heterogeneous and form cell-type-specific circuits with inhibitory interneurons^16–18^. Here, we found that the transcriptional regulator SATB2 plays an important role in CA1 PC differentiation to suppress synapse formation from PV+ interneurons to superficial PCs. This is necessary to establish differential strength of feedforward inhibition of Schaffer collateral input to superficial and deep PCs in CA1.

It has been hypothesized that PCs in CA1 have a laminar organization that is compressed within the stratum pyramidale^1,41^. Indeed, CA1 contains multiple PC subtypes that can be classified by projection target and molecular expression and are biased to either the deep or superficial sublayer^1,2,16^. The differentiation of these PCs and their radial position are correlated with their birth date^2,40,41^, but the molecular mechanisms remain obscure. Interestingly, CA1 PCs express the same set of transcription factors that are crucial for neocortical pyramidal cell differentiation, including *Satb2*, *Fezf2*, *Ctip2*, *Tbr1*, and *Sox5*^1,2,4,22,23,27,34,60–62^. However, previous work found that while *Sox5* and *Fezf2* are strongly biased to deep PCs, *Satb2* and *Ctip2* are expressed throughout the stratum pyramidale^2,34–36,63^. Thus, it was unclear if transcription factors characterized in neocortex contribute to PC differentiation in CA1. Our immunohistochemistry data are consistent with the observation that SATB2 is expressed in both deep and superficial PCs in juvenile mice. However, we found that during early postnatal development SATB2 is biased to later-born superficial PCs and progenitors. Thus, the relative timing of *Satb2* expression may be important for CA1 PC differentiation and contribute to the temporal fate-restriction of different subtypes^64–66^. Furthermore, although we observed SATB2 expression throughout CA1 after postnatal day 8, previous work found that *Satb2* transcription remains higher in superficial cells relative to deep in juvenile mice^2^. Thus, after the first postnatal week, differential *Satb2* transcription levels may also contribute to differences between deep and superficial PCs.

Previous work in adult mice found that deep PCs receive stronger feedforward inhibition of Schaffer collateral input relative to superficial PCs^19^. This is due to a higher density of perisomatic PV+ interneuron synapses surrounding deep PCs^16,17^, which results in larger unitary inhibitory synaptic currents^16^. We found that cKO of *Satb2* from PCs resulted in an increased number of SYT2+ inhibitory synaptic puncta surrounding superficial PCs. Importantly, the number of inhibitory synapses was not randomly dysregulated: in mutant mice, the number of synapses surrounding superficial PCs increased to match that observed around deep PCs, while the number of synapses surrounding deep PCs remained the same as in control mice. This suggests that loss of *Satb2* from superficial PCs altered their differentiation during development, causing them to resemble deep PCs during inhibitory synaptogenesis. In neocortex, proper PC differentiation also determines PV+ interneuron synapse formation. Ye et al. ^67^ found that IT-type PCs in layer 2/3 receive fewer PV+ synaptic puncta compared to layer 5 PT-type PCs. Strikingly, overexpressing *Fezf2* during embryonic development altered layer 2/3 IT-type PC differentiation, causing them to receive a comparable number of PV+ interneuron synaptic puncta as L5 PT-type PCs. Our data show that manipulating the expression of master transcriptional regulators in CA1 hippocampus causes inhibitory circuit changes like those observed for PC classes in neocortex, suggesting developmental mechanisms are conserved.

We found that deep PCs receive more PV+ interneuron synaptic puncta than superficial PCs as early as postnatal day 10. This suggests that differential feedforward inhibition between these cell-types is hardwired during development and dependent on PC cell-type identity. Indeed, in the absence of *Satb2*, the number of inhibitory synapses surrounding superficial PCs increased to equal deep PCs. How does *Satb2* expression in superficial PCs suppress PV+ interneuron synapse formation? It is likely that SATB2 downregulates expression of signaling molecules necessary to recruit or stabilize these synapses. Previous work in neocortex suggests likely candidate mechanisms. Wu et al. ^68^ found that layer 5 PT-type PCs express the chemokine ligand CXCL12 selectively at their soma to attract the axons of PV+ interneurons, which express the receptors CXCR4/CXCR7. Furthermore, they provide evidence that PT-type PCs use this mechanism to promote stronger inhibition relative to neighboring IT-type PCs^68,69^. In CA1, it is possible that *Satb2* expression in superficial PCs downregulates CXCL12 or a similar signaling molecule to reduce inhibitory synapse number relative to deep PCs. Alternatively, Favuzzi *et al.* ^55^ found that *Lgi2* expression is enriched in neocortical PV+ interneurons, and that knockdown results in reduced number of SYT2+ synapses around PC somas. They further show that neocortical PC somas express the LGI2 binding partner ADAM22 as a candidate postsynaptic molecular mechanism. PCs in CA1 also express ADAM22, thus, *Satb2* expression in superficial PCs may regulate its expression to control inhibitory synapse formation. However, ADAM22 also interacts with LGI1 to regulate excitatory synaptic strength from Schaffer collaterals^70^. We did not observe changes at this synapse in PC *Satb2* cKO mice, thus, other members of the ADAM family may be more likely candidates to explain our results^71^. Future work that focuses on cell-type-specific synaptic proteomics in CA1 will be necessary to determine the *Satb2*-dependent molecular mechanisms that regulate PV+ inhibitory synapses.

CA1 also contains a separate population of perisomatic-targeting basket cells that can be identified by expression of cholecystokinin (CCK)^72^. Valero *et al.* ^17^ found evidence using immunohistochemistry that CCK+ interneurons preferentially target superficial PCs, the opposite of PV+ interneurons. However, Lee *et al.* ^16^ did not find evidence for this using paired whole-cell recordings. Subsequently, Hartzell *et al.* ^18^ found that the strength of CCK+ interneuron synapses to superficial PCs, but not deep PCs, is plastic and activity-dependent. Specifically, when animals were exposed to an enriched environment, CCK+ interneurons formed additional synapses selectively to superficial PCs to enhance inhibition. This may explain the discrepancy between the previous studies. Interestingly, environmental enrichment drove expression of the immediate early gene NPAS4 in both superficial and deep PCs^18^. However, NPAS4 induced superficial PCs, but not deep PCs, to release an unknown retrograde signal to recruit CCK+ interneuron axon terminals^18^. It is tempting to speculate that SATB2 differentially regulates the targets of NPAS4 to regulate cell-type-dependent plasticity at CCK+ interneuron to superficial PC synapses.

In summary, we show that SATB2 regulates PC differentiation and inhibitory circuit development in CA1 hippocampus. Ultimately, it may play distinct roles in PCs further defined by a combination of their birth date, clonal lineage, and long-range axonal projection target^16,40,41^. Revealing the molecular mechanisms of CA1 PC differentiation will allow different cell-types to be classified and experimentally manipulated to understand hippocampal circuit function.

## STAR METHODS

KEY RESOURCES TABLE.

KEY RESOURCES TABLE
| REAGENT or RESOURCE | SOURCE | IDENTIFIER |
| --- | --- | --- |
| Antibodies |  |  |
| Mouse anti-NeuN | Abcam | Cat #: ab104224, RRID: AB_10711040 |
| Rabbit anti-NeuN | Abcam | Cat #: ab177487, RRID: AB_2532109 |
| Rabbit anti-PV | Abcam | Cat #: ab181086, RRID: AB_2924658 |
| Rabbit anti-SATB2 | Abcam | Cat #: ab92446, RRID: AB_10563678 |
| Mouse anti-SYT2 | Abcam | Cat #: ab154035, RRID: AB_2916272 |
| Goat anti-rabbit Alexa Fluor 488 | Thermo Fisher Scientific | Cat #: A-11008, RRID: AB_143165 |
| Goat anti-mouse Alexa Fluor 488 | Thermo Fisher Scientific | Cat #: A-11001, RRID: AB_2534069 |
| Goat anti-rabbit Alexa Fluor 555 | Thermo Fisher Scientific | Cat #: A-21428, RRID: AB_2535849 |
| Goat anti-mouse Alexa Fluor 555 | Thermo Fisher Scientific | Cat #: A-21422, RRID: AB_2535844 |
| DAPI | Thermo Fisher Scientific | Cat #: D1306 |
| Streptavidin Alexa Fluor 488 | Thermo Fisher Scientific | Cat #: S32354, RRID: AB_2315383 |
| Other Reagents |  |  |
| Biocytin | Sigma-Aldrich | Cat #: B4261 |
| CGP55845 | Sigma-Aldrich | Cat #: SML0594 |
| SR95531 | Sigma-Aldrich | Cat #: S106 |
| Experimental Models: Organisms/Strains |  |  |
| Mouse: Emx1<sup>IRES-Cre</sup> | The Jackson Laboratory | JAX: 005628 |
| Mouse: loxP flanked Satb2 | Rudolf Grosschedl (Srinivasan *et al.* <sup>33</sup>) | N/A |
| Mouse: Parvalbumin-TdTomato | The Jackson Laboratory | JAX: 027395 |
| Software and Algorithms |  |  |
| Igor Pro | WaveMetrics | N/A |
| pClamp 11.1 | Molecular Devices | N/A |
| NeuroMatic | Rothman and Silver <sup>73</sup> | N/A |
| LAS X | Leica Microsystems | N/A |
| Simple Neurite Tracer for Fiji | Longair *et al.* <sup>74</sup>, Schindelin *et al.* <sup>75</sup> | N/A |
| GraphPad | Prism | v. 10.1.1 |

## CONTACT FOR REAGENT AND RESOURCE SHARING

Further information and requests for resources and reagents should be directed to and will be fulfilled by the Lead Contact, Jason Wester.

## EXPERIMENTAL MODEL AND SUBJECT DETAILS

### Animals

All experiments were conducted in accordance with animal protocols approved by the Ohio State University IACUC. Both male and female mice were used without bias. Emx1^IRES-Cre^ mice were obtained from the Jackson Laboratory (stock no. 005628). Satb2^lox/lox^ (Srinivasan *et al.* ^33^) mice were obtained from the lab of Dr. Chris McBain (National Institutes of Health) with the permission of Dr. Rudolf Grosschedl (Max Planck Institute). Parvalbumin-TdTomato mice were obtained from the Jackson Laboratory (stock no. 027395). Immunohistochemistry, recording of basic membrane properties, and extracellular stimulation experiments were performed in Emx1^IRES-Cre^;Satb2^lox/lox^ mice. Paired whole-cell patch clamp recordings were performed in Emx1^IRES-Cre^;Satb2^lox/lox^;Parvalbumin-TdTomato mice.

## METHOD DETAILS

### Slice preparation

Juvenile mice (P21-P30) were anesthetized with isoflurane and then decapitated. The brain was dissected in ice-cold artificial cerebrospinal fluid (ACSF) containing (in mM): 100 sucrose, 80 NaCl, 3.5 KCl, 24 NAHCO3, 1.25 NaH2PO4, 4.5 MgCl, 0.5 CaCl2, and 10 glucose, saturated with 95% O_2_ and 5% CO_2_. Coronal slices of dorsal CA1 hippocampus (300 μm) were cut using a Leica VT 1200S vibratome (Leica Microsystems) and incubated in the above solution at 35 °C for 30 minutes post-dissection. Slices were then maintained at room temperature until use in recording ACSF containing (in mM): 130 NaCl, 3.5 KCl, 24 NaHCO3, 1.25 NaH2PO4, 1.5 MgCl, 2.5 CaCl2, and 10 glucose, saturated with 95% O_2_ and 5% CO_2_.

### Slice electrophysiology

For recording, slices were constantly perfused with ACSF at 2 mL/min at a temperature of 31-33 °C. Slices and cells were visualized using an upright microscope (Scientifica SliceScope) with a 40x water-immersion objective (Olympus), camera (Scientifica SciCam Pro), and Oculus software. Deep and superficial neurons were identified by their location within the pyramidal cell layer; superficial cells closest to stratum radiatum and deep cells closest to stratum oriens. TdTomato-expressing parvalbumin interneurons were identified by fluorescence (CoolLED pE-300ultra). Recording pipettes were pulled from borosilicate glass (World Precision Instruments) to a resistance of 3-5 MΩ using a vertical pipette puller (Narishige PC-100). Whole-cell patch clamp recordings were amplified using a Multiclamp 700B amplifier (Molecular Devices), filtered at 3 kHz (Bessel filter), and digitized at 20 kHz (Digidata 1550B and pClamp v11.1, Molecular Devices). Recordings were not corrected for liquid junction potential. Series resistance (10-25 MΩ) was closely monitored throughout recordings. Recordings were discarded if series resistance passed 25 MΩ. In current-clamp mode, cells were biased to a membrane potential (Vm) of –70 mV; in voltage-clamp mode, a holding potential of –70 mV was applied. The internal solution used to collect intrinsic membrane properties and excitatory postsynaptic currents and potentials contained (in mM): 130 K-gluconate, 5 KCl, 2 NaCl, 4 MgATP, 0.3 NaGTP, 10 phosphocreatine, 10 HEPES, 0.5 EGTA, and 0.2% biocytin. The calculated ECl-for this solution was –79 mV. pH was adjusted to 7.4 with KOH. The internal solution used to record GABA_A_-mediated currents in voltage-clamp at a holding potential of –70 mV contained (in mM): 85 K-gluconate, 45 KCl, 2 NaCl, 4 MgATP, 0.3 NaGTP, 10 phosphocreatine, 10 HEPES, and 0.5 EGTA, for a calculated ECl-of –29 mV. pH was adjusted to 7.4 with KOH.

Extracellular stimulation of Schaffer collaterals in the stratum radiatum and perforant path in the stratum lacunosum moleculare was performed with a constant current isolator (World Precision Instruments A365 Stimulus Isolator) to deliver current through a low resistance (∼750 kΩ) ACSF-filled glass pipette. Pulses were 0.1 ms long and given every 20 seconds, with three pulses per stimulus intensity. For stimulation of Schaffer collaterals, a cut was made during dissection at CA3/CA2 to prevent epileptic activity. For experiments in which GABA_A_ and GABA_B_ receptors were blocked, 2 μM SR95531 and 1 μM CGP55845 were added to the ACSF. For paired recordings of synaptic connections, presynaptic cells were made to fire trains of action potentials (25 at 50 Hz) using 2 nA steps of 2 ms duration every 10 s for 10 trials. Post-synaptic cells were recorded in voltage-clamp at –70 mV.

### Electrophysiology data analysis

All data were analyzed in Igor Pro (WaveMetrics) using custom routines. pClamp files were imported into Igor using NeuroMatic^73^. Input resistance (Rin) was measured using a linear regression of voltage deflections (±20 mV from –70 mV resting membrane potential) in response to 1 s current steps. To calculate voltage sag, the membrane potential was biased to –70 mV (V_initial) followed by injection of a 1 s negative current step of sufficient amplitude to reach a steady state Vm of –90 mV during the last 200 ms of the current injection (V_sag). The peak hyperpolarized Vm prior to sag (V_hyp) was used to calculate the sag index as (V_hyp – V_sag)/(V_hyp – V_initial); see Figure 2C. Capacitance was calculated as 1/Rin, both measured from the voltage response to a 10 pA current step relative to a –70 mV resting potential. Firing rate and adaptation ratio were collected from sweeps at twice threshold necessary to evoke an action potential. Adaptation ratio was calculated as the inter-spike interval between the first two action potentials divided by the averaged inter-spike interval between the last four action potentials.

To analyze postsynaptic currents in paired recordings, we first zeroed the data by subtracting the baseline and then performed 10 repetitions of binomial (Gaussian) smoothing. All postsynaptic currents (PSCs) were then analyzed relative to the timing of the peak of each presynaptic spike during the train. PSCs were detected by threshold crossing (–7 to –10 pA); if this threshold was already crossed at the instant of spike peak, the PSC data associated with that spike of that trial were discarded as being contaminated by spontaneous events. The proportion of failures was calculated as the ratio of evoked PSCs to the total number of non-contaminated trials for each presynaptic spike. The PSC potency was calculated as the average peak amplitude of all successfully evoked PSCs for each presynaptic spike (i.e., failures were not included in the average).

### Antibodies

Primary antibodies included: mouse anti-NeuN (1:500; Abcam, #ab104224), rabbit anti-Neun (1:500, Abcam, #ab177487), rabbit anti-PV (1:3000; Abcam, #ab181086), rabbit anti-SATB2 (1:100; Abcam, #ab92446), mouse anti-SYT2 (1:200; Abcam, #ab154035). DAPI was used at a concentration of 1:2000. Secondary antibodies were conjugated with Alexa Fluor dyes 488 or 555 (1:1000, Invitrogen).

### Immunohistochemistry

Mice were anesthetized with ketamine/xylazine and tissue was fixed via transcardial perfusion with 4% paraformaldehyde. Brains were post-fixed for 1 hour (age P21) or 4 hours (ages P10 and P14) in 4% paraformaldehyde. Brains were then cryopreserved in 30% sucrose. Sections were cut on a freezing microtome at 40 μm. Sections were rinsed in phosphate buffered saline Triton (PBS-Triton), blocked for 2 hours in blocking buffer consisting of 10% normal goat serum with 0.5% Triton X-100 and 1% BSA, and then incubated in primary antibody overnight at 4 °C. Sections were then rinsed with PBS and incubated in secondary antibody and DAPI for 2 hours at room temperature or overnight at 4 °C. All antibodies were diluted in carrier solution containing PBS with 1% BSA, 1% normal goat serum, and 0.5% Triton X-100. Sections were then rinsed, mounted on slides, and cover slipped.

### Recovery of cell morphology

Following slice electrophysiology, 300 μm tissue slices were drop-fixed for 24 hours in 4% paraformaldehyde. Slices were then rinsed with PBS and incubated in secondary antibody and DAPI overnight at 4 °C. All antibodies were diluted in carrier solution with PBS with 1% BSA, 1% normal goat serum, and 0.5% Triton X-100. Slices were next rinsed with PBS and resectioned on a freezing microtome at 100 μm and then slide mounted.

### Image acquisition and analysis

Biocytin filled cells were imaged from 100 μm thick hemisphere sections. Tissue for IHC was imaged from 40 μm thick sections. Sections were imaged using a Leica DM6 CS confocal microscope with a 10X (for PV in entire CA1), 20X (for biocytin-filled cells and SYT2 and SATB2 in entire CA1) or 63X (for SYT2+ puncta in stratum pyramidale) objective. Images were processed in Fiji^75^. For all fluorescence intensity quantifications, images were cropped to 200 μm wide. Borders of stratum pyramidale were determined by NeuN expression. The borders of strata oriens, radiatum, and lacunosum moleculare and the alveus were determined by DAPI expression. Within each cropped image, the fluorescence signal was normalized by setting the maximal intensity pixel to 255 and the minimal intensity pixel to 0. Within each image, a line profile analysis was run to average the pixels within each row. The line profile analyses were then averaged across multiple sections and multiple mice, with the center of stratum pyramidale (as determined by NeuN expression) as the alignment point. Finally, the intensity values were replotted on a range from 0-1. For manual cell and puncta counts, investigators were blinded to condition. Reconstructions were made using Simple Neurite Tracer^74^ for Fiji^75^.

## QUANTIFICATION AND STATISTICAL ANALYSIS

Distributions were first tested for normality using a Shapiro-Wilk test. Distributions that did not violate normality were compared using parametric tests, while those that did were compared using nonparametric tests. Statistical significance was set at p < 0.05. Data in the text and graphs are reported as mean ± SEM. Statistical tests were performed using Graphpad Prism (Graphpad Software, San Diego, CA).

## Author contributions

M.A.H and J.C.W. designed the experiments. M.A.H., N.B., A.S., D.N., A.H.M., A.C.J., and J.C.W. collected the data. M.A.H. and J.C.W. analyzed the data. M.A.H. and J.C.W. wrote the paper.

## Acknowledgements

This work was supported by funding from the NIH (R01MH124870) to J.C.W. We thank Chris McBain (NIH) for providing us with floxed Satb2 mice with the permission of Rudolf Grosschedl (Max Planck Institute).

---

**Supplemental Figure 1.**
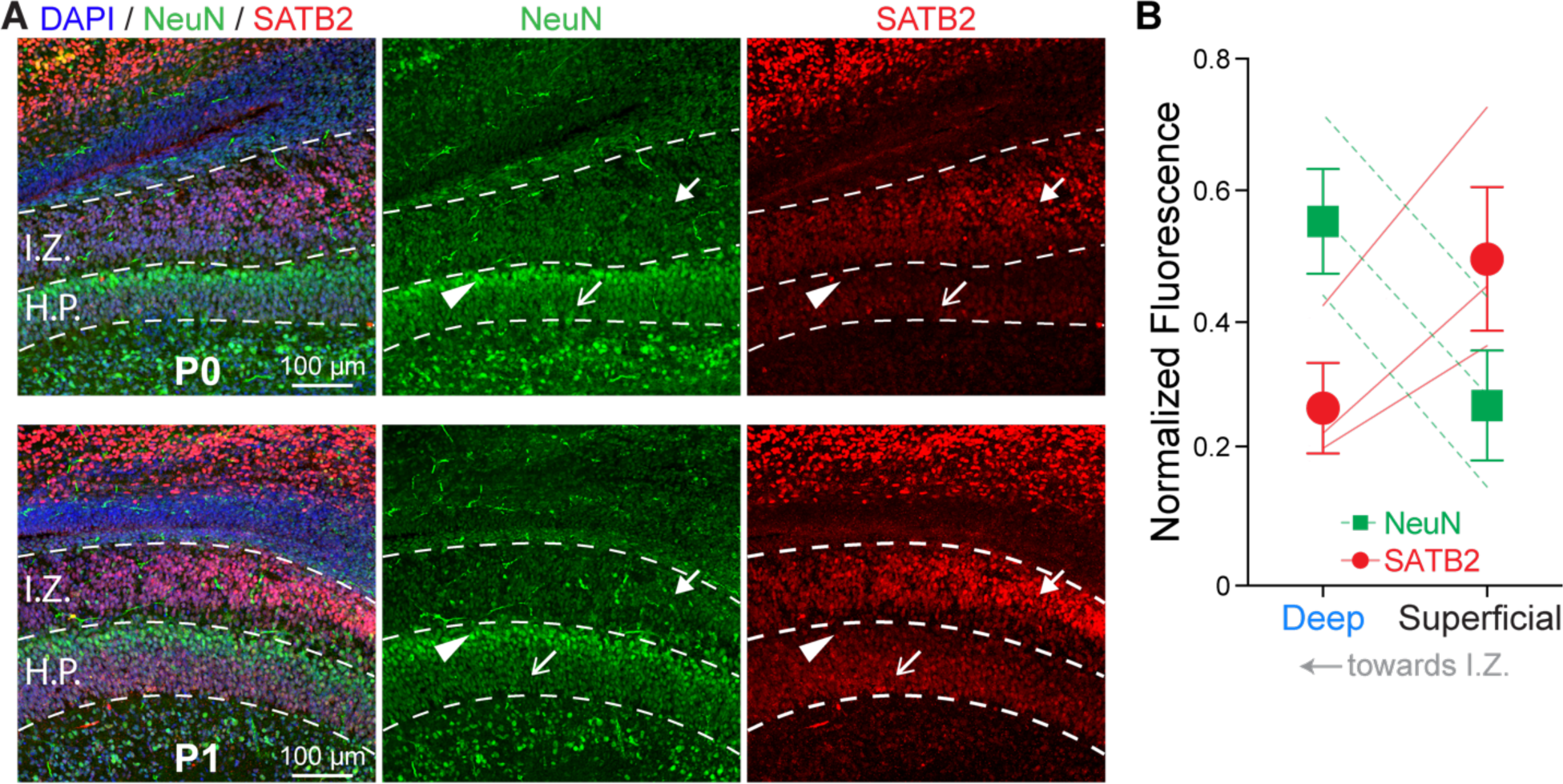
Related to Figure 1. SATB2 is preferentially expressed in superficial PCs and later-born progenitors during early postnatal development. (A) SATB2 expression in progenitor cells in the hippocampal intermediate zone (IZ) and early pyramidal cells in the hippocampal plate (HP) at postnatal days 0 and 1. Arrow: SATB2 expression in later-born migrating progenitor cells. Carrot: lack of SATB2 expression in early-born deep PCs. Arrow outline: SATB2 expression in later-born superficial PCs. (B) Quantification of NeuN and SATB2 normalized fluorescence in deep and superficial PCs at postnatal days 0, 1, and 2. For analysis, HP was split in half horizontally. The half closer to the IZ was labeled deep while the half further away was labeled superficial.

**Supplemental Figure 2.**
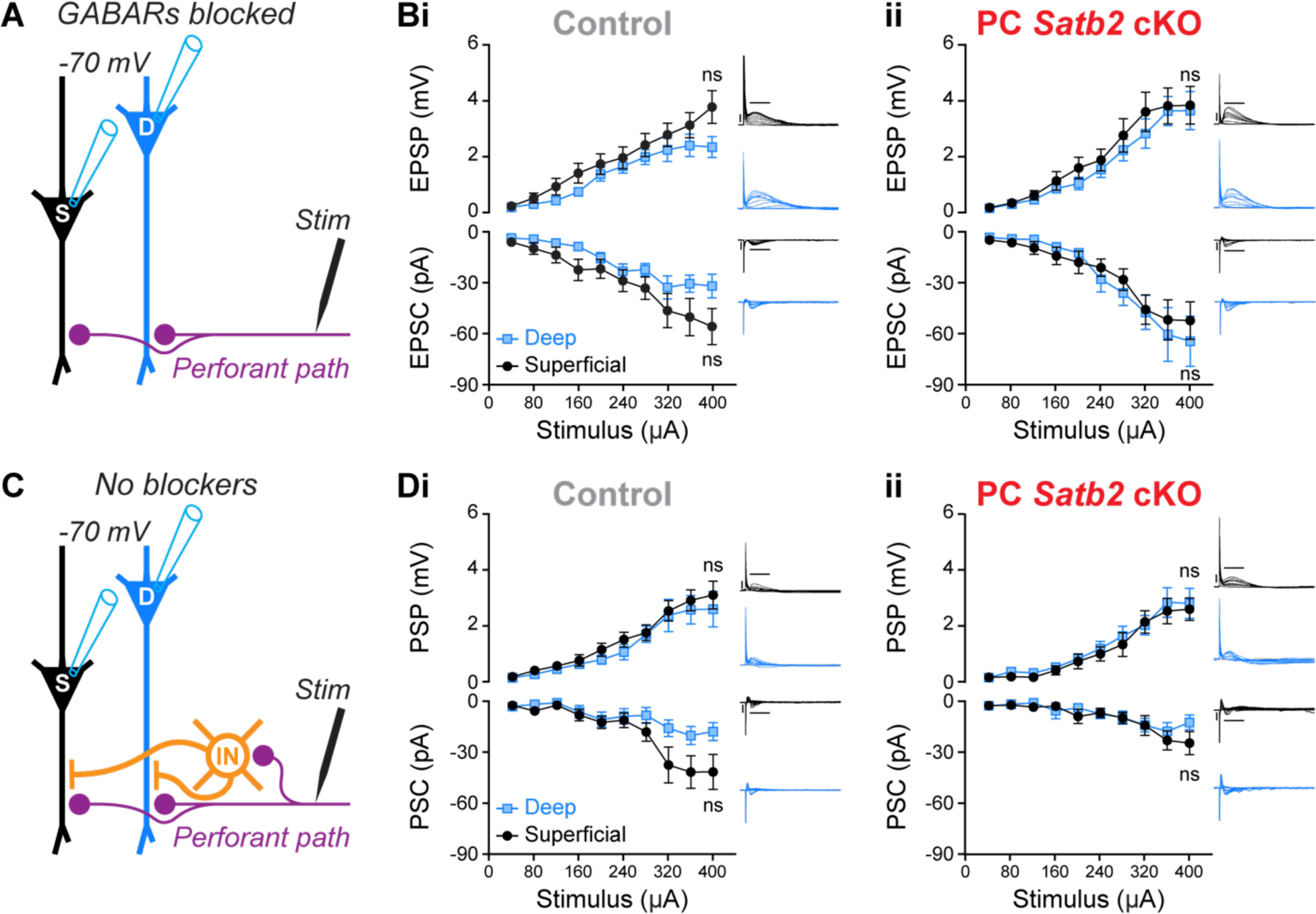
PC *Satb2* cKO does not impact perforant path input or feedforward inhibition of deep and superficial PCs. (A) Experimental configuration with inhibition blocked. (B) Input-output curves of PC responses to perforant path stimulation with inhibition blocked. PCs recorded in current-clamp (top) and voltage-clamp (bottom). (i) Control. Two-way ANOVA of EPSP (F[9, 140] = 0.6820, p = 0.72) and EPSC (F[9, 140] = 0.6803, p = 0.73). n = 8 deep and 8 superficial PCs from 7 mice. (ii) PC *Satb2* cKO. Two-way ANOVA of EPSP (F[9, 140] = 0.1710, p = 0.99) and EPSC (F[9, 140] = 0.3698, p = 0.95). n = 8 deep and 8 superficial PCs from 4 mice. Insets are example responses from single cells. Scale bars are 4 mV, 100 pA, 20 ms. (C) Experimental configuration with inhibition intact. (D) Same as (B) with inhibition intact. (i) Control. Two-way ANOVA of PSP (F[9, 140] = 0.1424, p = 0.99) and PSC (F[9, 140] = 1.765, p = 0.08). n = 8 deep and 8 superficial PCs from 8 mice. (ii) PC *Satb2* cKO. Two-way ANOVA of PSP (F[9, 130] = 0.09248, p = 0.99) and PSC (F[9, 140] = 0.6907, p = 0.71). n = 8 deep and 8 superficial PCs from 6 mice. Insets are example responses from single cells. Scale bars same as (B).

**Supplemental Figure 3.**
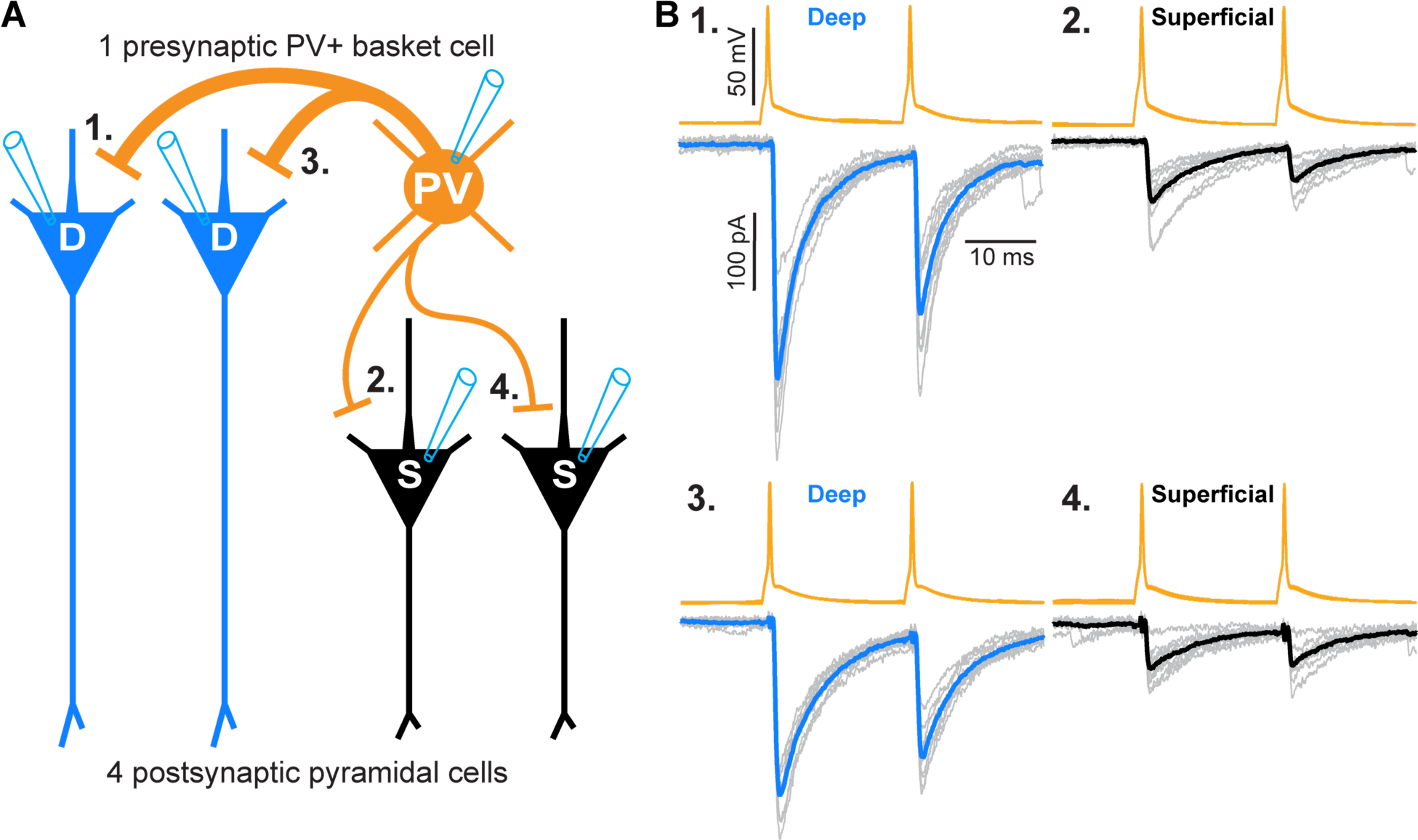
A single PV interneuron provides stronger synaptic input to deep PCs than superficial PCs. (A) Recording configuration. The four PCs were recorded in the order numbered in the figure, alternating deep and superficial. (B) Physiology of synaptic connections from the single PV interneuron to four PCs. Blue and black traces are averages of 10 single trials (gray) for each synaptic connection.

**Supplemental Figure 4.**
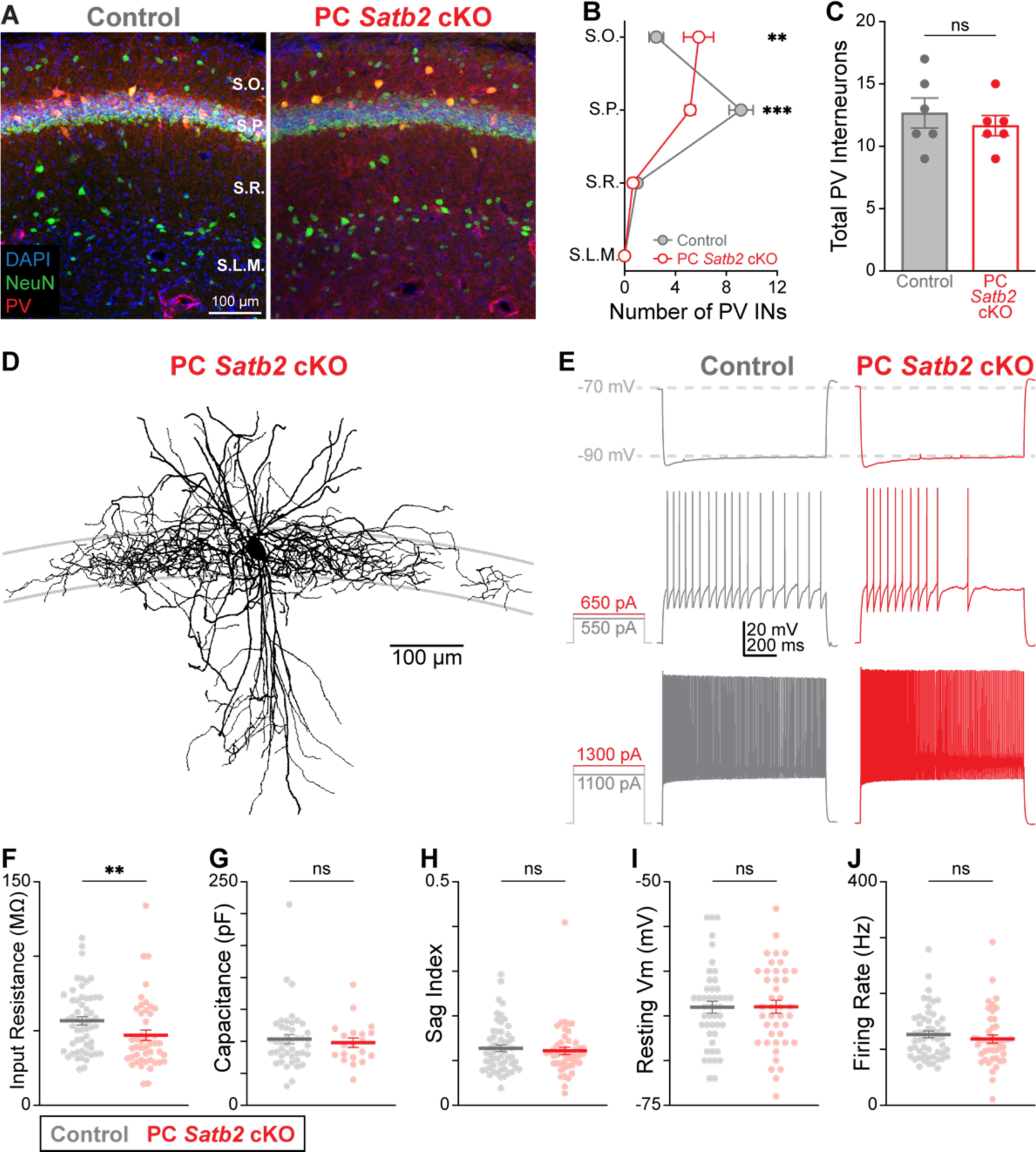
PC *Satb2* cKO disrupts PV interneuron migration but not physiology. (A) Examples of PV immunohistochemistry in CA1. (B) PC *Satb2* cKO results in mislamination of PV+ interneurons (n = 2 images per mouse, 3 mice per condition). ** p < 0.01, *** p < 0.001, two-way ANOVA (F[3, 40] = 12.35, p < 0.0001) with Šídák’s multiple comparisons test. (C) PC *Satb2* cKO does not alter the total number of PV+ interneurons (n = 2 images per mouse, 3 mice per condition). Unpaired t test (t(10) = 0.6919, p = 0.5047). (D) PV+ interneuron morphology is unaltered by PC *Satb2* cKO. (E) Sample PV+ interneuron physiology, highlighting voltage sag (top), firing patterns at threshold (middle) and 2X threshold (bottom) from control and PC *Satb2* cKO mice. (F) Input Resistance. Control n = 56 cells; PC *Satb2* cKO n = 46 cells. Mann-Whitney U test (U = 884, ** p = 0.0063). (G) Capacitance. Control n = 43 cells; PC *Satb2* cKO n = 20 cells. Mann-Whitney U test (U = 406, p = 0.7308). (H) Voltage sag index. Control n = 56 cells; PC *Satb2* cKO n = 47 cells. Mann-Whitney U test (U = 1266, p = 0.7404). (I) Resting membrane potential. Control n = 56 cells); PC *Satb2* cKO n = 47 cells. Mann-Whitney U test (U = 943.5, p = 0.8529) (J) Firing rate at 2X spike threshold. Control n = 53 cells; PC *Satb2* cKO n = 45 cells. Mann-Whitney U test (U = 1060, p = 0.3473). For (F) – (J), data are from 13 control and 12 mutant mice.

## REFERENCES

1. Slomianka, L., Amrein, I., Knuesel, I., Sorensen, J.C., and Wolfer, D.P. (2011). Hippocampal pyramidal cells: the reemergence of cortical lamination. Brain Struct Funct 216, 301–317. 10.1007/s00429-011-0322-0.

2. Cembrowski, M.S., Bachman, J.L., Wang, L., Sugino, K., Shields, B.C., and Spruston, N. (2016). Spatial Gene-Expression Gradients Underlie Prominent Heterogeneity of CA1 Pyramidal Neurons. Neuron 13, 01086–01087.

3. Cembrowski, M.S., and Spruston, N. (2019). Heterogeneity within classical cell types is the rule: lessons from hippocampal pyramidal neurons. Nat Rev Neurosci 20, 193–204. 10.1038/s41583-019-0125-5.

4. Soltesz, I., and Losonczy, A. (2018). CA1 pyramidal cell diversity enabling parallel information processing in the hippocampus. Nat Neurosci 21, 484–493.

5. Masurkar, A.V., Tian, C., Warren, R., Reyes, I., Lowes, D.C., Brann, D.H., and Siegelbaum, S.A. (2020). Postsynaptic integrative properties of dorsal CA1 pyramidal neuron subpopulations. J Neurophysiol 123, 980–992. 10.1152/jn.00397.2019.

6. Dong, H.W., Swanson, L.W., Chen, L., Fanselow, M.S., and Toga, A.W. (2009). Genomic-anatomic evidence for distinct functional domains in hippocampal field CA1. Proc Natl Acad Sci U S A 106, 11794–11799. 10.1073/pnas.0812608106.

7. Mizuseki, K., Diba, K., Pastalkova, E., and Buzsaki, G. (2011). Hippocampal CA1 pyramidal cells form functionally distinct sublayers. Nat Neurosci 14, 1174–1181. 10.1038/nn.2894.

8. Danielson, N.B., Zaremba, J.D., Kaifosh, P., Bowler, J., Ladow, M., and Losonczy, A. (2016). Sublayer-Specific Coding Dynamics during Spatial Navigation and Learning in Hippocampal Area CA1. Neuron 91, 652–665.

9. Fattahi, M., Sharif, F., Geiller, T., and Royer, S. (2018). Differential Representation of Landmark and Self-Motion Information along the CA1 Radial Axis: Self-Motion Generated Place Fields Shift toward Landmarks during Septal Inactivation. J Neurosci 38, 6766–6778. 10.1523/jneurosci.3211-17.2018.

10. Geiller, T., Fattahi, M., Choi, J.S., and Royer, S. (2017). Place cells are more strongly tied to landmarks in deep than in superficial CA1. Nature communications 8, 14531.

11. Sharif, F., Tayebi, B., Buzsáki, G., Royer, S., and Fernandez-Ruiz, A. (2021). Subcircuits of Deep and Superficial CA1 Place Cells Support Efficient Spatial Coding across Heterogeneous Environments. Neuron 109, 363–376.e366. 10.1016/j.neuron.2020.10.034.

12. Berndt, M., Trusel, M., Roberts, T.F., Pfeiffer, B.E., and Volk, L.J. (2023). Bidirectional synaptic changes in deep and superficial hippocampal neurons following in vivo activity. Neuron 111, 2984–2994.e2984. 10.1016/j.neuron.2023.08.014.

13. Senior, T.J., Huxter, J.R., Allen, K., O’Neill, J., and Csicsvari, J. (2008). Gamma oscillatory firing reveals distinct populations of pyramidal cells in the CA1 region of the hippocampus. J Neurosci 28, 2274–2286. 10.1523/jneurosci.4669-07.2008.

14. Navas-Olive, A., Valero, M., Jurado-Parras, T., de Salas-Quiroga, A., Averkin, R.G., Gambino, G., Cid, E., and de la Prida, L.M. (2020). Multimodal determinants of phase-locked dynamics across deep-superficial hippocampal sublayers during theta oscillations. Nature communications 11, 2217. 10.1038/s41467-020-15840-6.

15. Harvey, R.E., Robinson, H.L., Liu, C., Oliva, A., and Fernandez-Ruiz, A. (2023). Hippocampo-cortical circuits for selective memory encoding, routing, and replay. Neuron 111, 2076–2090.e2079. 10.1016/j.neuron.2023.04.015.

16. Lee, S.H., Marchionni, I., Bezaire, M., Varga, C., Danielson, N., Lovett-Barron, M., Losonczy, A., and Soltesz, I. (2014). Parvalbumin-Positive Basket Cells Differentiate among Hippocampal Pyramidal Cells. Neuron. 10.1016/j.neuron.2014.03.034.

17. Valero, M., Cid, E., Averkin, R.G., Aguilar, J., Sanchez-Aguilera, A., Viney, T.J., Gomez-Dominguez, D., Bellistri, E., and de la Prida, L.M. (2015). Determinants of different deep and superficial CA1 pyramidal cell dynamics during sharp-wave ripples. Nat Neurosci 18, 1281–1290.

18. Hartzell, A.L., Martyniuk, K.M., Brigidi, G.S., Heinz, D.A., Djaja, N.A., Payne, A., and Bloodgood, B.L. (2018). NPAS4 recruits CCK basket cell synapses and enhances cannabinoid-sensitive inhibition in the mouse hippocampus. Elife 7.

19. Masurkar, A.V., Srinivas, K.V., Brann, D.H., Warren, R., Lowes, D.C., and Siegelbaum, S.A. (2017). Medial and Lateral Entorhinal Cortex Differentially Excite Deep versus Superficial CA1 Pyramidal Neurons. Cell Rep 18, 148–160.

20. Greig, L.C., Woodworth, M.B., Galazo, M.J., Padmanabhan, H., and Macklis, J.D. (2013). Molecular logic of neocortical projection neuron specification, development and diversity. Nat Rev Neurosci. 10.1038/nrn3586.

21. Lodato, S., and Arlotta, P. (2015). Generating neuronal diversity in the mammalian cerebral cortex. Annu Rev Cell Dev Biol 31, 699–720.

22. Britanova, O., de Juan Romero, C., Cheung, A., Kwan, K.Y., Schwark, M., Gyorgy, A., Vogel, T., Akopov, S., Mitkovski, M., Agoston, D., et al. (2008). Satb2 is a postmitotic determinant for upper-layer neuron specification in the neocortex. Neuron 57, 378–392. 10.1016/j.neuron.2007.12.028.

23. Gyorgy, A.B., Szemes, M., de Juan Romero, C., Tarabykin, V., and Agoston, D.V. (2008). SATB2 interacts with chromatin-remodeling molecules in differentiating cortical neurons. Eur J Neurosci 27, 865–873. 10.1111/j.1460-9568.2008.06061.x.

24. McKenna, W.L., Ortiz-Londono, C.F., Mathew, T.K., Hoang, K., Katzman, S., and Chen, B. (2015). Mutual regulation between Satb2 and Fezf2 promotes subcerebral projection neuron identity in the developing cerebral cortex. Proc Natl Acad Sci U S A 112, 11702–11707.

25. Alcamo, E.A., Chirivella, L., Dautzenberg, M., Dobreva, G., Farinas, I., Grosschedl, R., and McConnell, S.K. (2008). Satb2 regulates callosal projection neuron identity in the developing cerebral cortex. Neuron 57, 364–377. 10.1016/j.neuron.2007.12.012.

26. Wester, J.C., Mahadevan, V., Rhodes, C.T., Calvigioni, D., Venkatesh, S., Maric, D., Hunt, S., Yuan, X., Zhang, Y., Petros, T.J., and McBain, C.J. (2019). Neocortical Projection Neurons Instruct Inhibitory Interneuron Circuit Development in a Lineage-Dependent Manner. Neuron 102, 960–975 e966. 10.1016/j.neuron.2019.03.036.

27. Chen, B., Schaevitz, L.R., and McConnell, S.K. (2005). Fezl regulates the differentiation and axon targeting of layer 5 subcortical projection neurons in cerebral cortex. Proc Natl Acad Sci U S A 102, 17184–17189.

28. Chen, B., Wang, S.S., Hattox, A.M., Rayburn, H., Nelson, S.B., and McConnell, S.K. (2008). The Fezf2-Ctip2 genetic pathway regulates the fate choice of subcortical projection neurons in the developing cerebral cortex. Proc Natl Acad Sci U S A 105, 11382–11387. 10.1073/pnas.0804918105.

29. Leone, D.P., Heavner, W.E., Ferenczi, E.A., Dobreva, G., Huguenard, J.R., Grosschedl, R., and McConnell, S.K. (2015). Satb2 Regulates the Differentiation of Both Callosal and Subcerebral Projection Neurons in the Developing Cerebral Cortex. Cereb Cortex 25, 3406–3419.

30. Lodato, S., Molyneaux, B.J., Zuccaro, E., Goff, L.A., Chen, H.H., Yuan, W., Meleski, A., Takahashi, E., Mahony, S., Rinn, J.L., et al. (2014). Gene co-regulation by Fezf2 selects neurotransmitter identity and connectivity of corticospinal neurons. Nat Neurosci 17, 1046–1054.

31. Lodato, S., Rouaux, C., Quast, K.B., Jantrachotechatchawan, C., Studer, M., Hensch, T.K., and Arlotta, P. (2011). Excitatory projection neuron subtypes control the distribution of local inhibitory interneurons in the cerebral cortex. Neuron 69, 763–779. 10.1016/j.neuron.2011.01.015.

32. Molyneaux, B.J., Arlotta, P., Hirata, T., Hibi, M., and Macklis, J.D. (2005). Fezl is required for the birth and specification of corticospinal motor neurons. Neuron 47, 817–831.

33. Srinivasan, K., Leone, D.P., Bateson, R.K., Dobreva, G., Kohwi, Y., Kohwi-Shigematsu, T., Grosschedl, R., and McConnell, S.K. (2012). A network of genetic repression and derepression specifies projection fates in the developing neocortex. Proc Natl Acad Sci U S A 109, 19071–19078.

34. Nielsen, J.V., Blom, J.B., Noraberg, J., and Jensen, N.A. (2010). Zbtb20-induced CA1 pyramidal neuron development and area enlargement in the cerebral midline cortex of mice. Cereb Cortex 20, 1904–1914. 10.1093/cercor/bhp261.

35. Sugiyama, T., Osumi, N., and Katsuyama, Y. (2014). A novel cell migratory zone in the developing hippocampal formation. J Comp Neurol 522, 3520–3538. 10.1002/cne.23621.

36. Simon, R., Brylka, H., Schwegler, H., Venkataramanappa, S., Andratschke, J., Wiegreffe, C., Liu, P., Fuchs, E., Jenkins, N.A., Copeland, N.G., et al. (2012). A dual function of Bcl11b/Ctip2 in hippocampal neurogenesis. The EMBO journal 31, 2922–2936.

37. Zhang, L., Song, N.N., Zhang, Q., Mei, W.Y., He, C.H., Ma, P., Huang, Y., Chen, J.Y., Mao, B., Lang, B., and Ding, Y.Q. (2019). Satb2 is required for the regionalization of retrosplenial cortex. Cell Death Differ. 10.1038/s41418-019-0443-1.

38. Angevine, J.B., Jr. (1965). Time of neuron origin in the hippocampal region. An autoradiographic study in the mouse. Exp Neurol Suppl 2, 1–70.

39. Bayer, S.A. (1980). Development of the hippocampal region in the rat. I. Neurogenesis examined with 3H-thymidine autoradiography. J Comp Neurol 190, 87–114. 10.1002/cne.901900107.

40. Cavalieri, D., Angelova, A., Islah, A., Lopez, C., Bocchio, M., Bollmann, Y., Baude, A., and Cossart, R. (2021). CA1 pyramidal cell diversity is rooted in the time of neurogenesis. Elife 10. 10.7554/eLife.69270.

41. Huszár, R., Zhang, Y., Blockus, H., and Buzsáki, G. (2022). Preconfigured dynamics in the hippocampus are guided by embryonic birthdate and rate of neurogenesis. Nat Neurosci 25, 1201–1212. 10.1038/s41593-022-01138-x.

42. Marissal, T., Bonifazi, P., Picardo, M.A., Nardou, R., Petit, L.F., Baude, A., Fishell, G.J., Ben-Ari, Y., and Cossart, R. (2012). Pioneer glutamatergic cells develop into a morpho-functionally distinct population in the juvenile CA3 hippocampus. Nature communications 3, 1316. 10.1038/ncomms2318.

43. Altman, J., and Bayer, S.A. (1990). Prolonged sojourn of developing pyramidal cells in the intermediate zone of the hippocampus and their settling in the stratum pyramidale. J Comp Neurol 301, 343–364. 10.1002/cne.903010303.

44. Gorski, J.A., Talley, T., Qiu, M., Puelles, L., Rubenstein, J.L., and Jones, K.R. (2002). Cortical excitatory neurons and glia, but not GABAergic neurons, are produced in the Emx1-expressing lineage. J Neurosci 22, 6309–6314.

45. Kaiser, T., Ting, J.T., Monteiro, P., and Feng, G. (2016). Transgenic labeling of parvalbumin-expressing neurons with tdTomato. Neuroscience 321, 236–245. 10.1016/j.neuroscience.2015.08.036.

46. Harb, K., Magrinelli, E., Nicolas, C.S., Lukianets, N., Frangeul, L., Pietri, M., Sun, T., Sandoz, G., Grammont, F., Jabaudon, D., et al. (2016). Area-specific development of distinct projection neuron subclasses is regulated by postnatal epigenetic modifications. Elife 5, e09531.

47. Hattox, A.M., and Nelson, S.B. (2007). Layer V neurons in mouse cortex projecting to different targets have distinct physiological properties. J Neurophysiol 98, 3330–3340. 10.1152/jn.00397.2007.

48. Morishima, M., and Kawaguchi, Y. (2006). Recurrent connection patterns of corticostriatal pyramidal cells in frontal cortex. J Neurosci 26, 4394–4405. 10.1523/jneurosci.0252-06.2006.

49. Pla, R., Borrell, V., Flames, N., and Marin, O. (2006). Layer acquisition by cortical GABAergic interneurons is independent of Reelin signaling. J Neurosci 26, 6924–6934. 10.1523/jneurosci.0245-06.2006.

50. Hevner, R.F., Daza, R.A., Englund, C., Kohtz, J., and Fink, A. (2004). Postnatal shifts of interneuron position in the neocortex of normal and reeler mice: evidence for inward radial migration. Neuroscience 124, 605–618. 10.1016/j.neuroscience.2003.11.033.

51. Stevens, C.F., and Wang, Y. (1995). Facilitation and depression at single central synapses. Neuron 14, 795–802. 10.1016/0896-6273(95)90223-6.

52. Allen, C., and Stevens, C.F. (1994). An evaluation of causes for unreliability of synaptic transmission. Proc Natl Acad Sci U S A 91, 10380–10383. 10.1073/pnas.91.22.10380.

53. Sommeijer, J.P., and Levelt, C.N. (2012). Synaptotagmin-2 is a reliable marker for parvalbumin positive inhibitory boutons in the mouse visual cortex. PLoS One 7, e35323. 10.1371/journal.pone.0035323.

54. Danglot, L., Triller, A., and Marty, S. (2006). The development of hippocampal interneurons in rodents. Hippocampus 16, 1032–1060. 10.1002/hipo.20225.

55. Favuzzi, E., Deogracias, R., Marques-Smith, A., Maeso, P., Jezequel, J., Exposito-Alonso, D., Balia, M., Kroon, T., Hinojosa, A.J., E, F.M., and Rico, B. (2019). Distinct molecular programs regulate synapse specificity in cortical inhibitory circuits. Science 363, 413–417. 10.1126/science.aau8977.

56. Rozenberg, F., Robain, O., Jardin, L., and Ben-Ari, Y. (1989). Distribution of GABAergic neurons in late fetal and early postnatal rat hippocampus. Brain Res Dev Brain Res 50, 177–187. 10.1016/0165-3806(89)90193-4.

57. Nitsch, R., Bergmann, I., Kuppers, K., Mueller, G., and Frotscher, M. (1990). Late appearance of parvalbumin-immunoreactivity in the development of GABAergic neurons in the rat hippocampus. Neurosci Lett 118, 147–150.

58. de Lecea, L., del Rio, J.A., and Soriano, E. (1995). Developmental expression of parvalbumin mRNA in the cerebral cortex and hippocampus of the rat. Brain Res Mol Brain Res 32, 1–13.

59. Pouille, F., and Scanziani, M. (2004). Routing of spike series by dynamic circuits in the hippocampus. Nature 429, 717–723. 10.1038/nature02615.

60. Li, Y., You, Q.L., Zhang, S.R., Huang, W.Y., Zou, W.J., Jie, W., Li, S.J., Liu, J.H., Lv, C.Y., Cong, J., et al. (2017). Satb2 Ablation Impairs Hippocampus-Based Long-Term Spatial Memory and Short-Term Working Memory and Immediate Early Genes (IEGs)-Mediated Hippocampal Synaptic Plasticity. Mol Neurobiol. 10.1007/s12035-017-0531-5.

61. Jaitner, C., Reddy, C., Abentung, A., Whittle, N., Rieder, D., Delekate, A., Korte, M., Jain, G., Fischer, A., Sananbenesi, F., et al. (2016). Satb2 determines miRNA expression and long-term memory in the adult central nervous system. Elife 5.

62. Nielsen, J.V., Thomassen, M., Mollgard, K., Noraberg, J., and Jensen, N.A. (2014). Zbtb20 defines a hippocampal neuronal identity through direct repression of genes that control projection neuron development in the isocortex. Cereb Cortex 24, 1216–1229. 10.1093/cercor/bhs400.

63. Xie, Z., Ma, X., Ji, W., Zhou, G., Lu, Y., Xiang, Z., Wang, Y.X., Zhang, L., Hu, Y., Ding, Y.Q., and Zhang, W.J. (2010). Zbtb20 is essential for the specification of CA1 field identity in the developing hippocampus. Proc Natl Acad Sci U S A 107, 6510–6515. 10.1073/pnas.0912315107.

64. Walsh, C., and Cepko, C.L. (1993). Clonal dispersion in proliferative layers of developing cerebral cortex. Nature 362, 632–635. 10.1038/362632a0.

65. Desai, A.R., and McConnell, S.K. (2000). Progressive restriction in fate potential by neural progenitors during cerebral cortical development. Development (Cambridge, England) 127, 2863–2872.

66. Frantz, G.D., and McConnell, S.K. (1996). Restriction of late cerebral cortical progenitors to an upper-layer fate. Neuron 17, 55–61.

67. Ye, Z., Mostajo-Radji, M.A., Brown, J.R., Rouaux, C., Tomassy, G.S., Hensch, T.K., and Arlotta, P. (2015). Instructing Perisomatic Inhibition by Direct Lineage Reprogramming of Neocortical Projection Neurons. Neuron 88, 475–483.

68. Wu, P.R., Cho, K.K., Vogt, D., Sohal, V.S., and Rubenstein, J.L. (2016). The Cytokine CXCL12 Promotes Basket Interneuron Inhibitory Synapses in the Medial Prefrontal Cortex. Cereb Cortex, 6.

69. Lee, A.T., Gee, S.M., Vogt, D., Patel, T., Rubenstein, J.L., and Sohal, V.S. (2014). Pyramidal neurons in prefrontal cortex receive subtype-specific forms of excitation and inhibition. Neuron 81, 61–68. 10.1016/j.neuron.2013.10.031.

70. Lovero, K.L., Fukata, Y., Granger, A.J., Fukata, M., and Nicoll, R.A. (2015). The LGI1-ADAM22 protein complex directs synapse maturation through regulation of PSD-95 function. Proc Natl Acad Sci U S A 112, E4129–4137. 10.1073/pnas.1511910112.

71. Kegel, L., Aunin, E., Meijer, D., and Bermingham, J.R. (2013). LGI proteins in the nervous system. ASN Neuro 5, 167–181. 10.1042/an20120095.

72. Hefft, S., and Jonas, P. (2005). Asynchronous GABA release generates long-lasting inhibition at a hippocampal interneuron-principal neuron synapse. Nat Neurosci 8, 1319–1328. 10.1038/nn1542.

73. Rothman, J.S., and Silver, R.A. (2018). NeuroMatic: An Integrated Open-Source Software Toolkit for Acquisition, Analysis and Simulation of Electrophysiological Data. Front Neuroinform 12, 14.

74. Longair, M.H., Baker, D.A., and Armstrong, J.D. (2011). Simple Neurite Tracer: open source software for reconstruction, visualization and analysis of neuronal processes. Bioinformatics 27, 2453–2454. 10.1093/bioinformatics/btr390.

75. Schindelin, J., Arganda-Carreras, I., Frise, E., Kaynig, V., Longair, M., Pietzsch, T., Preibisch, S., Rueden, C., Saalfeld, S., Schmid, B., et al. (2012). Fiji: an open-source platform for biological-image analysis. Nature methods 9, 676–682.

